# A comprehensive foundation model for cryo-EM image processing

**DOI:** 10.1101/2024.11.04.621604

**Authors:** Yang Yan, Shiqi Fan, Fajie Yuan, Huaizong Shen

**Affiliations:** Research Center for Industries of The Future (RCIF), Westlake University, Hangzhou, Zhejiang, China; School of Engineering, Westlake University, Hangzhou, Zhejiang, China; Zhejiang Key Laboratory of Structural Biology, School of Life Sciences, Westlake University; Hangzhou, Zhejiang, China; Westlake Laboratory of Life Sciences and Biomedicine, Hangzhou, Zhejiang, China; Institute of Biology, Westlake Institute for Advanced Study, Hangzhou, Zhejiang, China

## Abstract

Cryogenic electron microscopy (cryo-EM) has become a premier technique for high-resolution structural determination of biological macromolecules^1–4^. However, its widespread adoption is hampered by the need for specialized expertise. We introduce the Cryo-EM Image Evaluation Foundation (Cryo-IEF) model, pre-trained on an extensive dataset of approximately 65 million cryo-EM particle images using unsupervised learning. Cryo-IEF excels in various cryo-EM data processing tasks, such as classifying particles from different structures, clustering particles by pose, and assessing the quality of particle images. Upon fine-tuning, the model effectively ranks particle images by quality at high efficiency, enabling the creation of CryoWizard—a fully automated single-particle cryo-EM data processing pipeline. This pipeline has successfully resolved high-resolution structures of diverse properties and proven adept at mitigating the prevalent preferred orientation challenge in many cryo-EM samples. The Cryo-IEF model and CryoWizard pipeline collectively represent a significant advancement in rendering cryo-EM technology more accessible, efficient, and robust, with substantial implications for life sciences research.

## Main

Cryogenic electron microscopy (cryo-EM) has become an indispensable tool for the structural analysis of biological macromolecules^1–4^. Unlike crystallography, cryo-EM does not require crystallization, which is often a significant obstacle for many biological specimens^5^. The past decade has witnessed a transformative leap in cryo-EM^6–8^, particularly in single-particle analysis, driven by advancements in direct detector devices and computational methods^9–14^, establishing it as a leading technique for structural elucidation.

Despite its potential, the broader application of cryo-EM is limited by the intricate data processing required, posing a substantial barrier to non-experts^15,16^. A typical cryo-EM workflow involves multiple steps, each traditionally requiring manual oversight, which can be labor-intensive, time-consuming, and prone to human error, potentially leading to suboptimal outcomes. To address these challenges, deep learning-based methods have been developed to automate specific workflow segments^17–23^. However, these solutions tend to be task-specific and still require considerable user intervention. Moreover, challenges such as structural heterogeneity^21,24^ and preferred orientation^25–27^ continue to complicate structure determination from many biological samples, despite numerous attempted remedies.

Pre-trained vision foundation models have proven extremely effective for a variety of image processing tasks, often outperforming traditional training methods on specialized datasets^28–31^. These models represent a new standard in image processing, with broad applications across various domains, including medical image analysis^32–35^ and the processing of fluorescence microscopy images^36^. Within the realm of cryo-EM data processing, several pivotal steps, such as particle pose estimation and 2D/3D classification, could potentially benefit from a pre-trained foundation model tailored to particle images. However, to the best of our knowledge, no such foundation models have yet been introduced.

In this study, we introduce the cryo-EM Image Evaluation Foundation Model (Cryo-IEF), tailored for a broad range of cryo-EM image processing tasks. The model demonstrates superior performance in the classification of particles from different structures, the clustering of particles by pose, and the assessment of particle quality. Building on this foundation model, we have developed CryoWizard, a fully automated pipeline for cryo-EM data processing, which further streamlines the cryo-EM workflow.

### An overview of Cryo-IEF

Cryo-IEF employs a contrastive learning framework akin to MoCo v3 to learn feature representations from cryo-EM particle images (Fig. 1)^30^. Each particle image in the pre-training dataset is processed to create two separate, independently augmented views, which are then encoded using two parallel encoders. A variety of data augmentation techniques are employed, such as random cropping, color jittering, blurring, solarization, and rotation. The core objective during training is to enhance the similarity of feature representations between different views of the same particle image while reducing similarity between views of distinct particle images. For detailed model architecture descriptions and training processes, please refer to the Methods (Extended Data Fig. 6a,b).

**Fig. 1.**
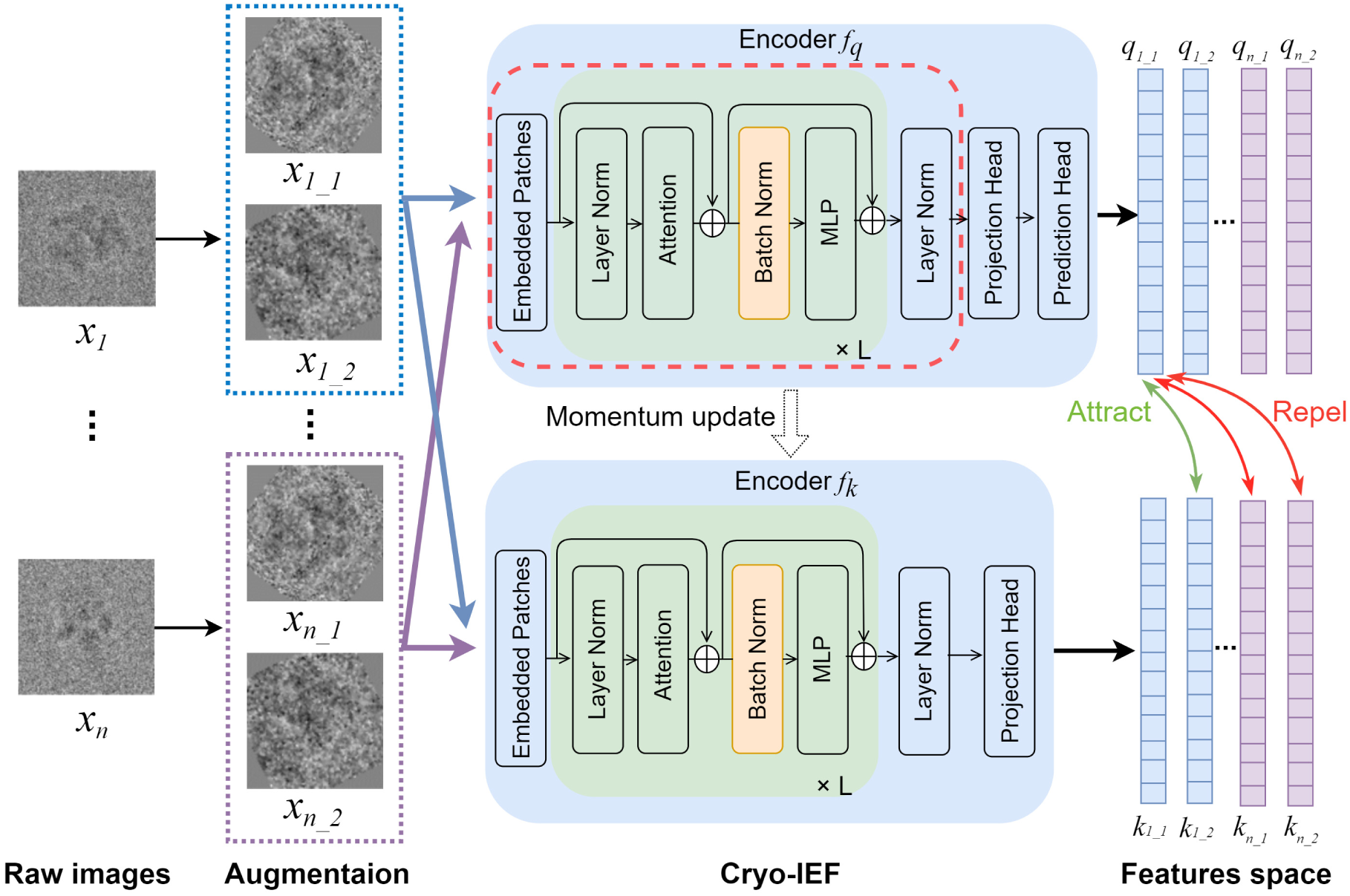
Pre-training framework of Cryo-IEF. Cryo-IEF utilizes a contrastive learning framework for pre-training. Two distinctly augmented views are generated from each particle image of the training datasets and encoded by two parallel encoders. The model is trained to maximize the similarity between the two views of the same image while minimizing the similarity between views from different particle images. Only the modules within the red dashed box are employed for downstream cryo-EM image processing tasks. For detailed information on the model architecture, please refer to the Methods.

The foundation model was trained using an extensive dataset comprising approximately 65 million cryo-EM particle images, which included over a hundred distinct biological structures. This diverse collection of datasets was amassed from several sources: the Electron Microscopy Public Image Archive (EMPIAR) database^37^, a previous study^38^, and our own in-house experimental data, with comprehensive details provided in the Methods section (Extended Data Fig. 7 and Extended Data Tables 1 and 2).

To rigorously assess the performance of the Cryo-IEF model across a broad spectrum of cryo-EM image processing tasks, we conducted semantic representation analysis on the Cryo-IEF-extracted features from test datasets not used in model training. These datasets include 12 simulated and 4 genuine particle datasets (Tables 1 and 2). The simulated datasets were generated from high-resolution cryo-EM maps downloaded from the Electron Microscopy Data Bank (EMDB) and were labeled with ground-truth information regarding structural variations and pose positions (Table 1). The molecular weights of the related structures range from 69 kDa to 1 MDa for the simulated datasets and from 68 kDa to 2.2 MDa for the genuine datasets. The details of these test datasets can be found in the Methods.

**Table 1.** Test datasets of simulated cryo-EM particle images.

| Particle Category | EMDB ID (Overall Molecular Weight) |  |  |
| --- | --- | --- | --- |
| 0-200 kDa | EMD-27990<br>(69 kDa) | EMD-4608<br>(86 kDa) | EMD-9648<br>(195 kDa) |
| 200-600 kDa | EMD-24057<br>(218 kDa) | EMD-0097<br>(328 kDa) | EMD-26062<br>(449 kDa) |
| 600-2000 kDa | EMD-10847<br>(669 kDa) | EMD-16651<br>(1 MDa) | EMD-10102<br>(958 kDa) |
| With Nucleic<br>Acids | EMD-41856<br>(523 kDa) | EMD-29879<br>(484 kDa) | EMD-24823<br>(210 kDa) |

**Table 2.** Test datasets of genuine cryo-EM particle images.

| EMPIAR ID | Overall<br>Molecular<br>Weight<br>(kDa) | Raw pixel<br>size (Å) | Accelerating<br>Voltage<br>(kV) | Spherical<br>Aberration<br>(mm) | Total<br>exposure<br>dose<br>(e/Å<sup>2</sup>) |
| --- | --- | --- | --- | --- | --- |
| EMPIAR-10535<sup>†</sup> | 68 | 1.038 | 200 | 2.7 | 47.13 |
| EMPIAR-11043<sup>†</sup> | 774 | 0.830 | 300 | 2.7 | 58.20 |
| EMPIAR-11387<sup>†</sup> | 1313 | 0.968 | 200 | 2.7 | 40.00 |
| EMPIAR-11388<sup>†*</sup> | 1500, 2200 | 0.800 | 300 | 4.0 | 40.00 |
| EMPIAR-10556<sup>‡</sup> | 725 | 0.822 | 300 | 2.7 | 83.00 |
| EMPIAR-11292<sup>‡</sup> | 900 | 0.860 | 300 | 2.7 | 40.00 |
| EMPIAR-10405<sup>‡</sup> | 440 | 0.435 | 200 | 2.7 | 60.00 |
| EMPIAR-10250<sup>‡</sup> | 64 | 0.556 | 300 | 2.7 | 69.00 |
| EMPIAR-10876<sup>‡</sup> | 400 | 0.540 | 300 | 2.7 | 49.90 |
| EMPIAR-10217<sup>‡◇</sup> | 347 | 0.662 | 300 | 2.7 | 61.25 |
| EMPIAR-10096<sup>◇</sup> | 150 | 1.310 | 300 | 2.7 | 82.00 |

<sup>†</sup> Datasets for CryoRanker evaluation.930 <sup>‡</sup> Datasets for CryoWizard evaluation.931 <sup>◇</sup> Datasets for preferred orientation correction test.
\* This structure has multiple conformations.

### Cryo-IEF classifies particles from different structures

Efficient and reliable classification of particles from different structures is crucial in cryo-EM data processing^24^, as it not only allows the resolving of each structure, but also improves the throughput of structural determination, potentially enabling resolving multiple structures in one collected dataset.

Recent studies have demonstrated that many pre-trained vision foundation models possess three dimensional awareness^39^. To test whether Cryo-IEF also has such capability and classifies particles from different structures, we conducted an initial assessment using k-Neareast Neighbors (k-NN) classification on test dataset particles. The features extracted by Cryo-IEF from 12 simulated datasets and 4 genuine datasets exhibited distinct clustering behaviors based on structural variations, achieving k-NN accuracies of 96.42% and 93.32%, respectively (Fig. 2). These results demonstrate that Cryo-IEF effectively differentiates particles from distinct structures.

**Fig. 2.**
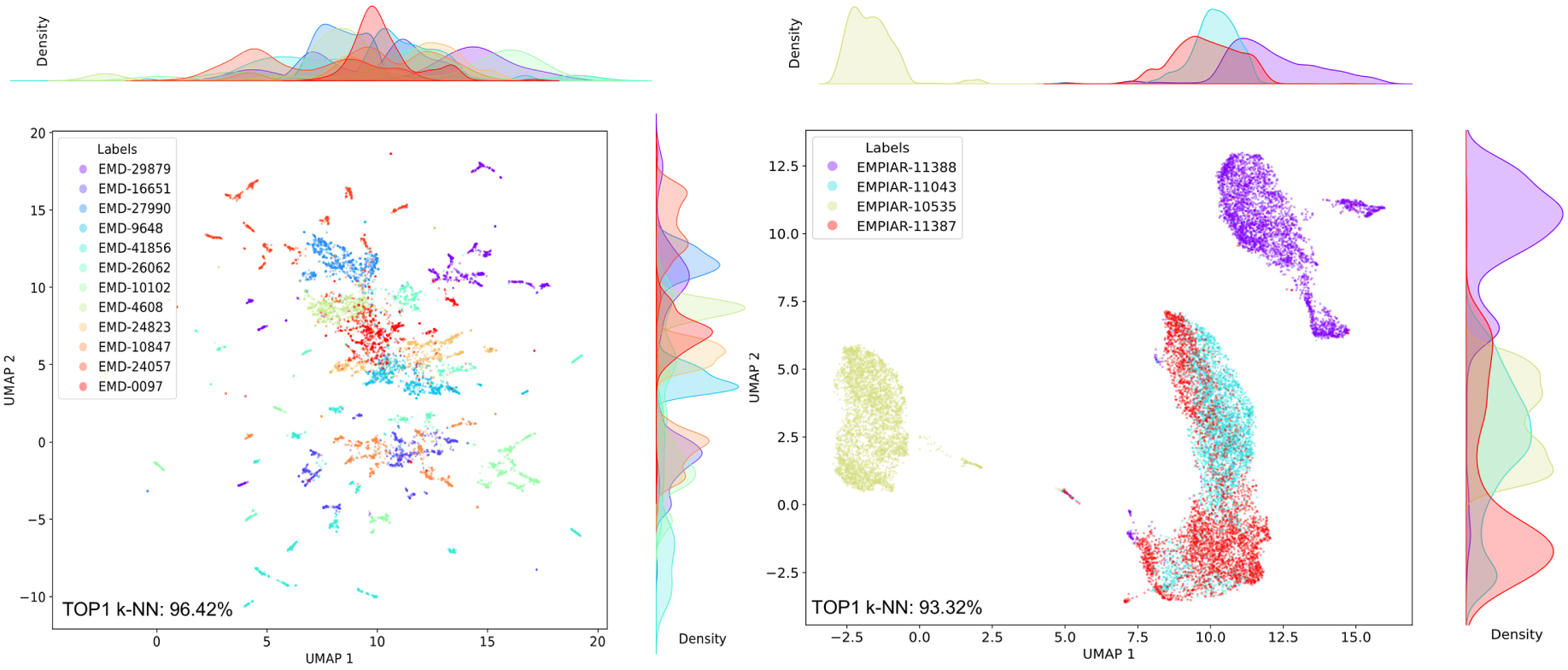
Cryo-IEF’s performance in classifying particles from different structures. Features extracted by Cryo-IEF from particles in 12 simulated datasets (left panel) and 4 genuine datasets (right panel) are visualized using Uniform Manifold Approximation and Projection (UMAP). These particle features are color-coded and labeled with their ground-truth EMDB or EMPIAR identifier numbers. The k-Nearest Neighbors (k-NN) scores for the particles from the simulated and genuine datasets are 96.42% and 93.32%, respectively, indicating that Cryo-IEF effectively classifies cryo-EM particles from various structures.

To further explore Cryo-IEF’s capability in classifying particles from different structures, we applied it to deep learning-based heterogeneous structural reconstruction tasks. Existing methods for such tasks often employ an autoencoder-like framework, where the structural reconstruction involves two main steps: an encoder projects particle images into a feature space, and a decoder reconstructs the 3D structure from this space^21,40–42^.

Typically, both the encoder and decoder are randomly initialized and trained together iteratively—the encoder to capture structural heterogeneity and the decoder to represent these structures in three dimensions.

In our study, we introduced a novel heterogeneous structural reconstruction AI model named CryoSolver (Fig. 3a). This model integrates Cryo-IEF as a fixed feature extractor within the DRGN-AI framework, replacing the standard trainable encoder^42^. Unlike traditional approaches, Cryo-IEF remains frozen with fixed parameters, focusing solely on feature extraction, while the decoder remains trainable. We evaluated CryoSolver using a resampled version of the Ribosembly dataset from CryoBench, which includes four distinct structures (Fig. 3b)^43^. The dataset features particles with varying proportions: 12.75% for structure 8C9C, 62.32% for 8C99, 16.82% for 8C93, and 8.11% for 8C8X. Detailed dataset information and training processes are provided in the Methods section. In constrast to the distribution of randomly initialized features without any clustering (Fig. 3c), the Cryo-IEF-extracted features from the test dataset particles were well-clustered, with particles from the same structure closely distributed (Fig. 3d). Following the hierarchical pose search process and stochastic gradient descent training of the decoder, we randomly selected three points for each structure from the feature space to perform 3D reconstruction. The reconstructions based on randomly initialized features predominantly resembled the second ground-truth structure (PDB ID: 8C99), which was the majority in the dataset (62.32%) (Fig. 3e). On the contrary, the structures generated by CryoSolver closely matched the ground-truth 3D structures (Fig. 3f). These findings indicate that Cryo-IEF, despite being trained in an unsupervised manner without explicit structural heterogeneity labels, effectively extracts features that reflect 3D structural variations, thereby demonstrating its capability to classify particles from different structures.

**Fig. 3.**
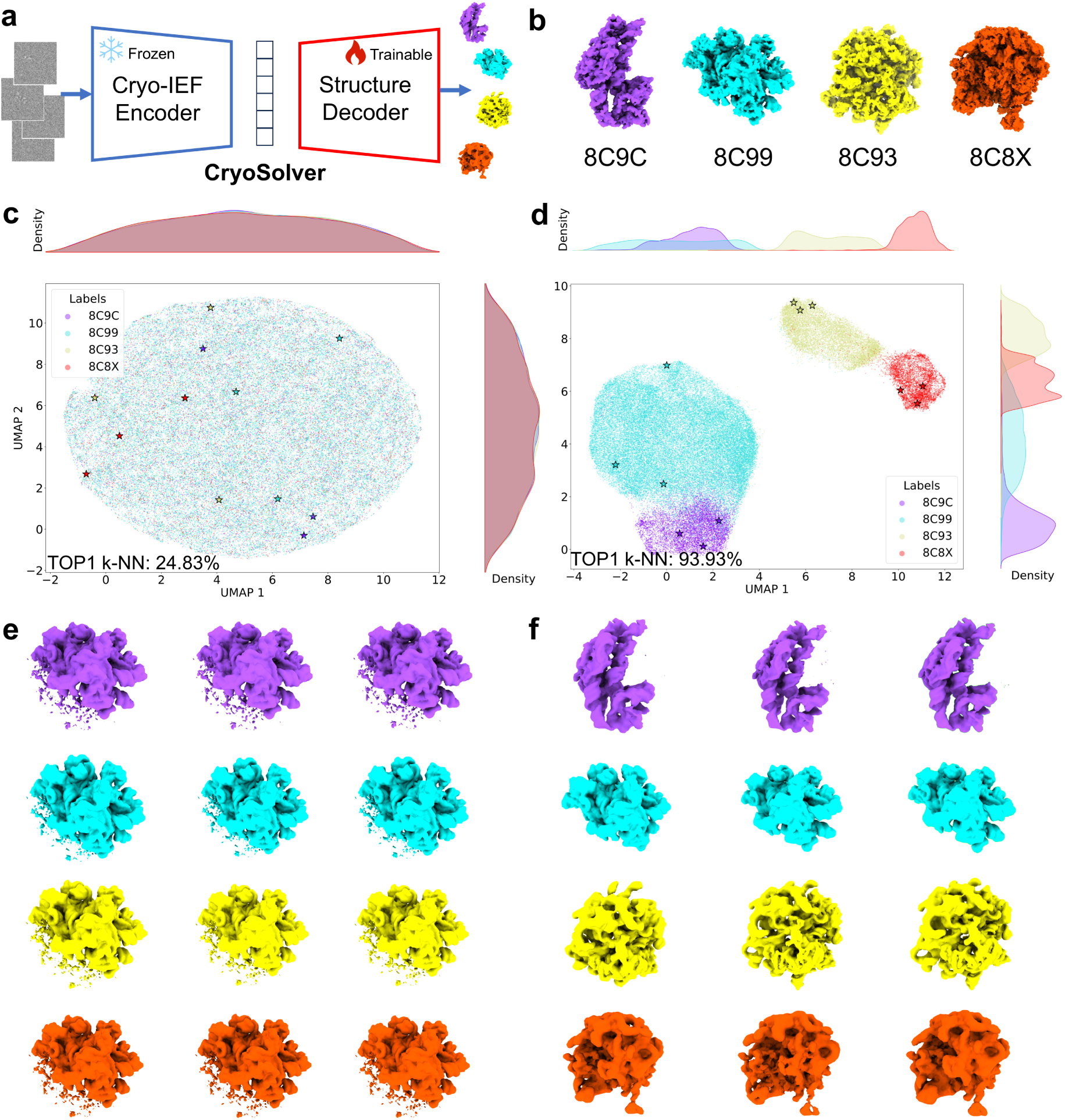
Reconstruction of heterogeneous structures from features extracted by Cryo-IEF. (**a**) The framework of CryoSolver for heterogeneous structural reconstruction. CryoSolver integrates Cryo-IEF with frozen parameters as the encoder and the trainable structure decoder of DRGN-AI for reconstructing heterogeneous structures. (**b**) Ground-truth structural maps corresponding to the test dataset are displayed, with their respective PDB codes labeled. (**c**)(**d**) Visualization of particle features from the test dataset, color-coded by their originating structures and displayed using UMAP. Panel (**c**) shows features initialized randomly, while panel (**d**) shows features extracted by Cryo-IEF. (**e**)(**f**) Reconstruction of individual structures from particle features marked with stars in panels (**c**) and (**d**) using the structure decoder. The reconstructed structures are shown in corresponding colors in panels (**e**) and (**f**), respectively.

### Cryo-IEF clusters particles by pose

In cryo-EM, the pose of a particle refers to its orientation and position within the context of its three-dimensional reconstruction. Accurate pose determination is crucial for achieving high-resolution cryo-EM maps^13,14,44^. Additionally, many cryo-EM samples suffer from the preferred orientation problem^25–27,45^, where most particles of a dataset adopt a limited number of dominant poses, leading to incomplete structural information. Accuratly determining the pose of each particle can help reduce these preferred orientations, leading to more accurate reconstructions.

The Cryo-IEF model is trained to maximize its ability to identify unique particle features, which suggests that Cryo-IEF can cluster particles with similar poses. To test this hypothesis, we evaluated the model’s performance in pose clustering using 12 simulated datasets, by employing the k-NN classification. Indeed, Cryo-IEF-extracted particle features demonstrated distinct clustering patterns that matched the ground-truth pose classes, achieving a mean k-NN accuracy of 90.6% (Fig. 4). The model showed exceptional performance for large molecular weight samples (over 400 kDa), with k-NN scores exceeding 95%. These results confirm that Cryo-IEF can effectively cluster cryo-EM particles by their poses. Further processing was conducted for the dataset EMD-24057. The particles of each cluster in EMD-24057 underwent 2D classification in CryoSPARC^14^, which revealed clear 2D class average images. Remarkably, the adjacent clusters visualized in the Uniform Manifold Approximation and Projection (UMAP)^46^ plot corresponded to particles with similar 2D class average images, indicating that the spatial distribution of features extracted by Cryo-IEF reflects the relationship between different particle poses, further demonstrating its effectiveness in pose clustering.

**Fig. 4.**
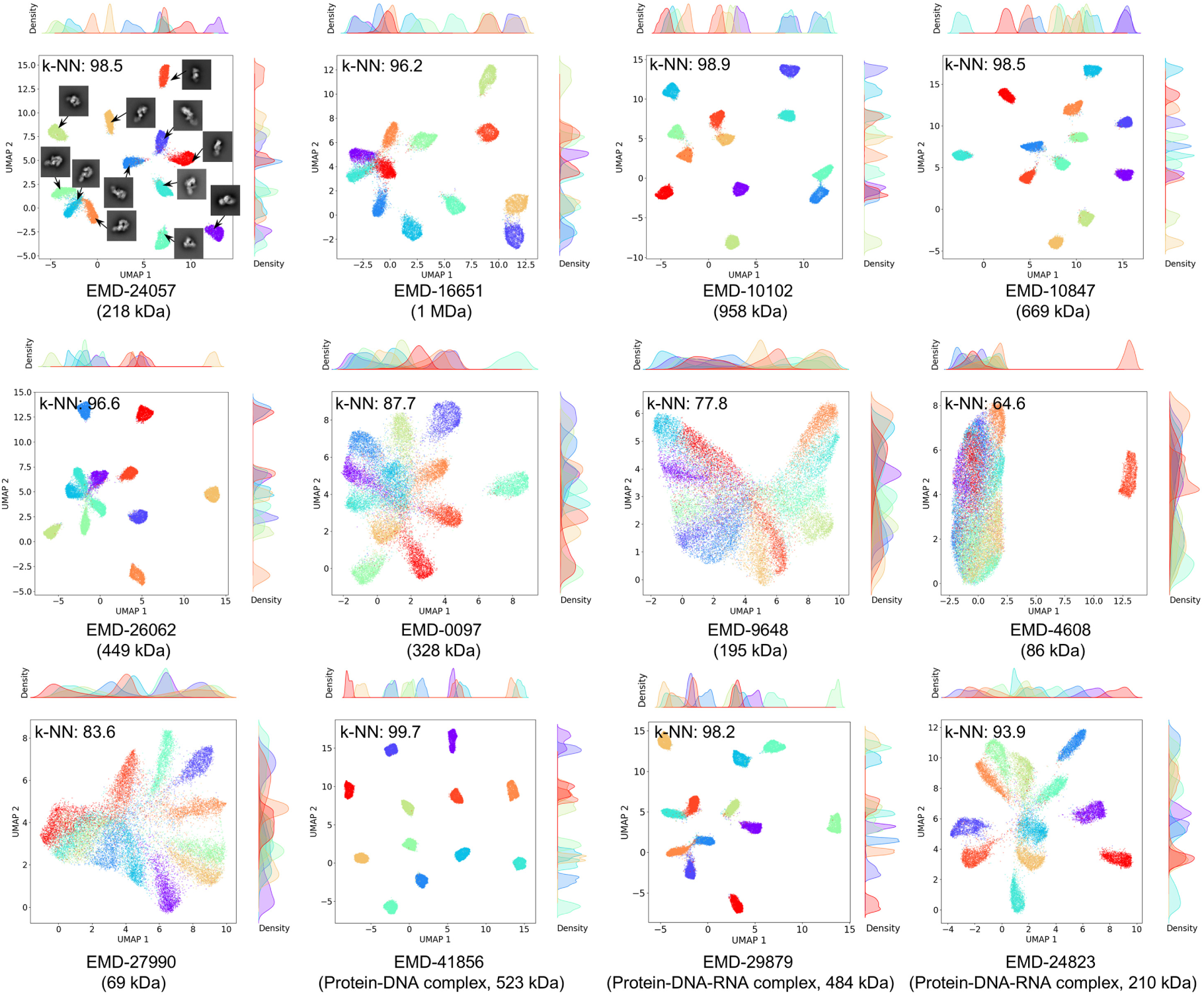
Cryo-IEF’s performance in clustering particles by pose. The particle features of the 12 simulated datasets, extracted by Cryo-IEF, are color-coded according to their corresponding ground-truth pose clusters and visualized with UMAP. The distribution of these particle features and the calculated k-NN scores demonstrate that Cryo-IEF effectively clusters cryo-EM particles according to their poses. For data generated from map EMD-24057, particles from each cluster were further processed using the 2D classification job in CryoSPARC. The resulting 2D class averages are displayed adjacent to their corresponding feature clusters in the panel.

### Cryo-IEF distinguishes particle quality

The primary objective of cryo-EM data processing is to achieve high-resolution structures by accurately aligning particle images^47^. This requires selectively retaining high-quality particles while discarding those that could compromise the resolution^15,44,48,49^. Here, particle quality is defined by the extent to which a particle contributes to the overall quality and resolution of the final constructed cryo-EM maps. Therefore, distinguishing and ranking particle images by quality is crucial for successful reconstruction. Cryo-EM images typically have extremely low signal-to-noise ratios (SNR), making it challenging to discern the image quality^50,51^. Consequently, Current algorithms cannot evaluate images individually; instead, particles with similar views are clustered through 2D classification to create averaged images, which are then collectively retained or discarded^13,14^. This approach, however, requires manual intervention and can introduce errors in decision-making on single images.

To tackle this challenge, we evaluated Cryo-IEF’s ability to assess cryo-EM image quality on a per-particle basis. We applied the k-NN algorithm to four genuine test datasets. Initially, particles underwent 2D classification in CryoSPARC, and each class was manually assigned a quality score based on the clarity of the class averages, ranging from 0 (pure noise or ice) to 1 (clear structural features) (Extended Data Fig. 1). Based on these scores, particles were categorized into high-quality (above 0.7), medium-quality (between 0.3 and 0.7), and low-quality (below 0.3) classes. Cryo-IEF-extracted features displayed distinct clustering with a k-NN accuracy of 74.06% (Fig. 5b), indicating that Cryo-IEF can differentiate particle quality without explicit training.

**Fig. 5.**
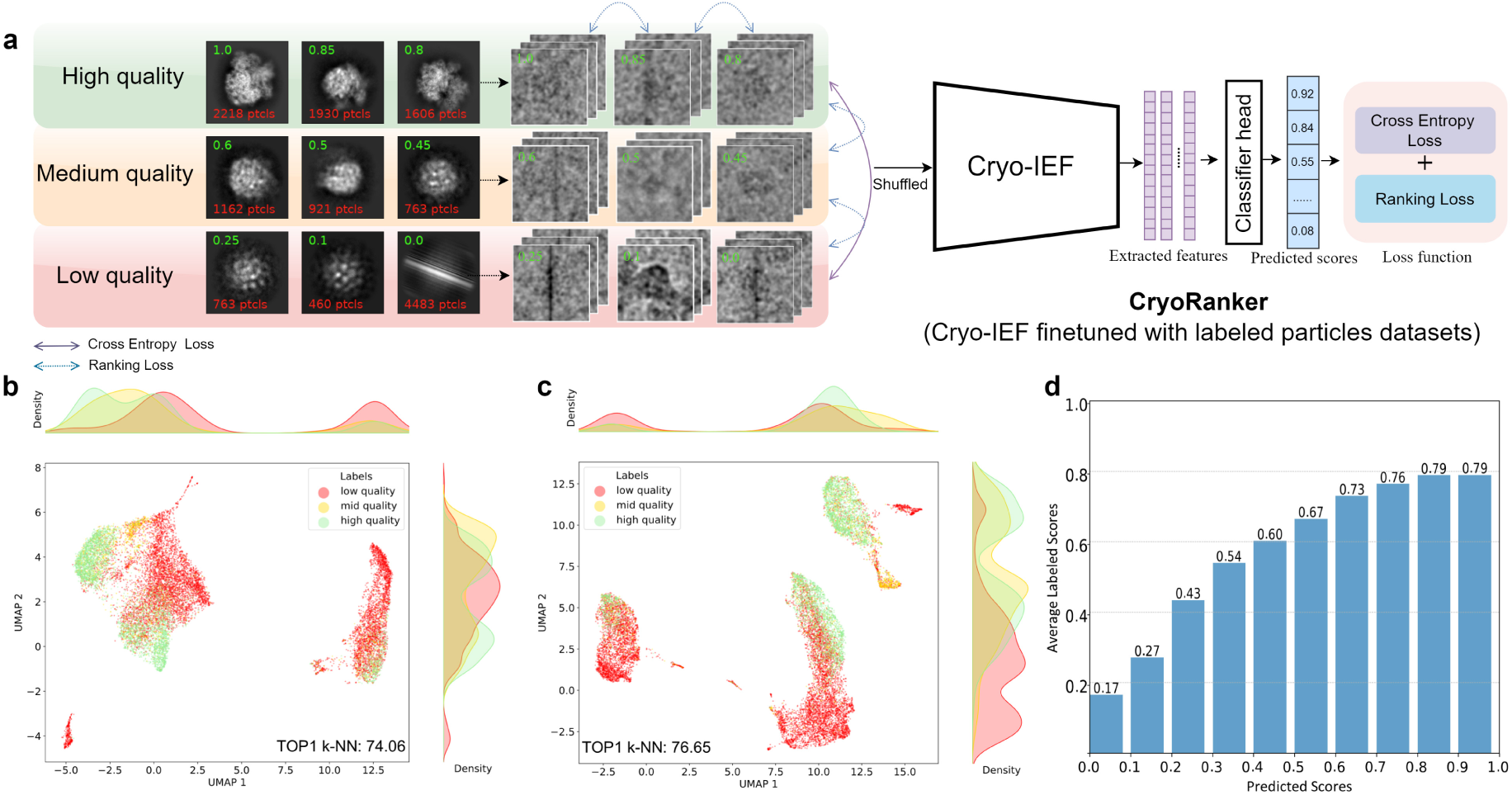
Cryo-IEF’s performance in ranking particles by quality. (**a**) Fine-tuning framework of CryoRanker. Cryo-IEF was fine-tuned to create CryoRanker, which ranks particles according to quality. During fine-tuning, particles were processed through CryoSPARC’s 2D classification job and manually assigned quality scores from 0 to 1, reflecting the quality of their corresponding 2D class averages. The optimization employed a combination of cross-entropy and margin ranking losses. Particles scoring above 0.7 were deemed high-quality, those below 0.3 low-quality, and scores falling in between indicated mid-quality. Further details on the quality score assignment are provided in Extended Data Fig. 1. (**b**)(**c**) Distributions of features extracted by Cryo-IEF (**b**) and CryoRanker (**c**) from four genuine datasets are color-coded according to their labeled qualities and visualized with UMAP. The results indicate that both models can distinguish particles of different qualities, with the fine-tuned CryoRanker achieving a higher k-NN score. (**d**) The distribution of the manually labeled average quality scores for test datasets, in relation to the predicted ones, reveals a positive correlation between the two sets of scores.

To improve Cryo-IEF’s capability in distinguishing and ranking particle quality, we developed CryoRanker, a fine-tuned version of Cryo-IEF, which incorporates Cryo-IEF’s backbone encoder with an additional classifier head (Fig. 5a and Extended Data Fig. 6c). A combination of cross-entropy and margin ranking losses were employed for fine-tuning. CryoRanker was trained on a comprehensive dataset of approximately 42 million labeled cryo-EM particle images, the quality scores of which were assigned using the same criteria as for the test datasets. Details of the training datasets and process are provided in the Methods section (Extended Data Fig. 7). Features extracted by CryoRanker from the four genuine test datasets showed clearer clustering with a k-NN score of 76.65%, surpassing Cryo-IEF’s performance (Fig. 5c). The predicted scores by CryoRanker closely matched the average labeled scores (Fig. 5d), supporting the ranking ability of the model. CryoRanker was also evaluated with precision and recall metrics, which reveal higher values for high- and low-quality particles than medium-quality ones (Extended Data Fig. 2). It is worth noting that CryoRanker exhibits high processing efficiency. In our tests, ranking one million particles required only an average of 35 minutes when using four NVIDIA V100 GPUs.

### An automated single-particle cryo-EM data processing pipeline

Multiple rounds of 2D and 3D classifications in cryo-EM data processing are resource-intensive, time-consuming, and often require human intervention. To streamline these processes, we developed CryoWizard, a fully automated cryo-EM data processing pipeline, by integrating CryoRanker to replace the 2D/3D classification steps (Fig. 6a and Extended Data Fig. 3). Currently, CryoWizard is designed only for homogeneous structure reconstruction.

**Fig. 6.**
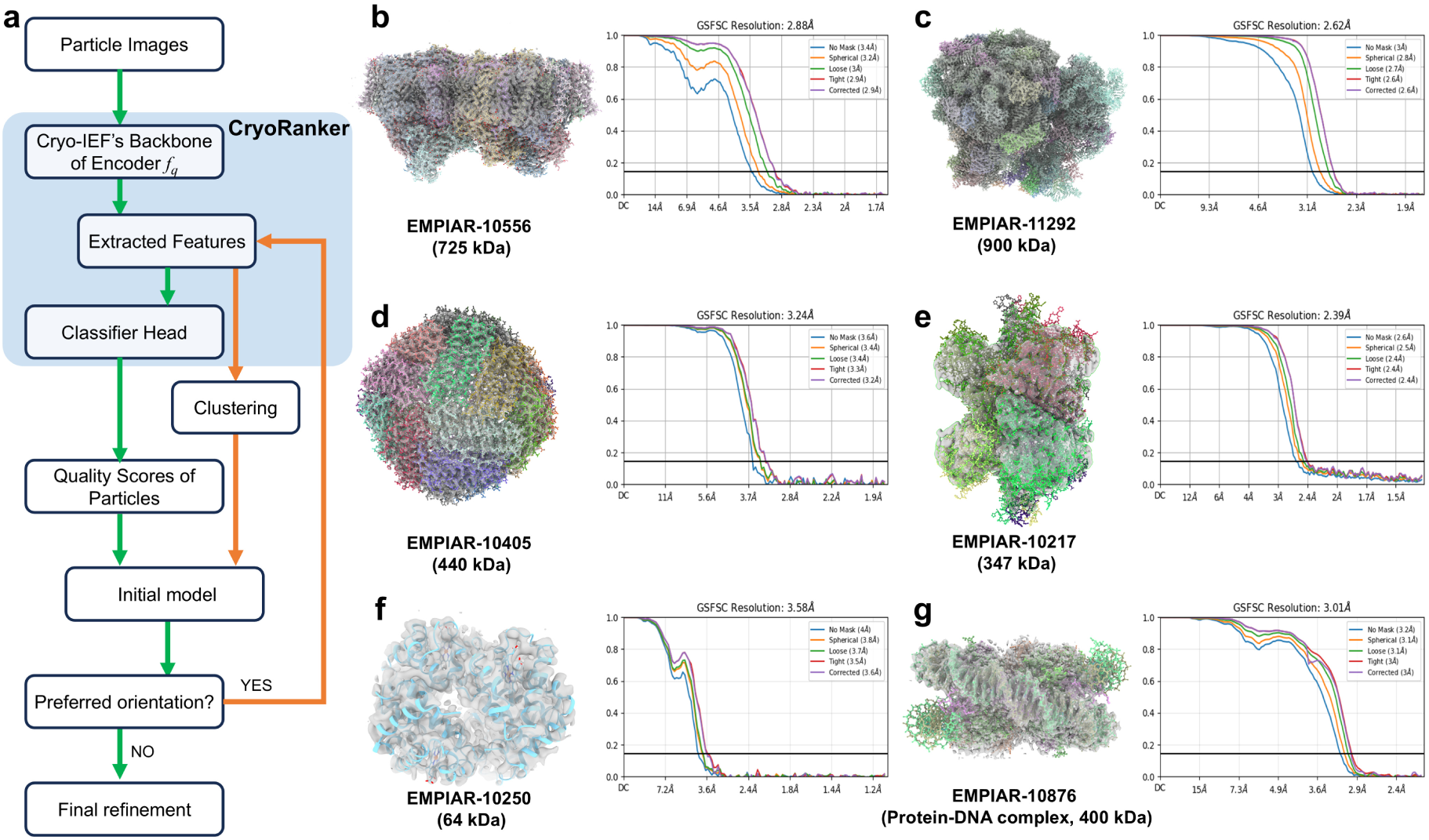
Performance of the fully automated cryo-EM data processing pipeline, CryoWizard. (**a**) A simplified data processing flowchart of CryoWizard. CryoWizard is a fully automated cryo-EM data processing pipeline with recorded cryo-EM movies/micrographs or particle images as inputs and the final maps as outputs. CryoRanker plays a central role in the pipeline by replacing the time- and resource-consuming, multiple rounds of 2D/3D classification jobs which requires human interventions. For details, please refer to the Methods and Extended Data Fig. 3. (**b-g**) Cryo-EM maps automatically resolved by the pipeline are aliganed with their respective PDB models. Fourier Shell Correlation (FSC) curves for each reconstruction are displayed to indicate the resolved resolutions. EMPIAR ID and molecular weights of the datasets are also labeled. The figures of aligned maps and models were prepared using ChimeraX^58^.

CryoWizard comprises three main stages which are data preprocessing and particle quality ranking, initial model reconstruction, and final map refinement (Fig. 6a and Extended Data Fig. 3). First, collected cryo-EM movies or micrographs are preprocessed to produce extracted particles, which are ranked and assigned quality scores by CryoRanker.

Second, optimal number of top-ranked particles (e.g., 50,000) are selected to generate an initial model, based on which a refined map is reconstructed as the template for downstream structural refinement. Third, a larger optimal number of top-ranked particles (e.g. 200,000) are chosen for the final high-resolution map refinement. For datasets afflicted by preferred orientation issues, an additional clustering module is implemented in the pipeline to address this problem, as detailed in the next section. Except for the assignment of quality scores by CryoRanker, the remaining tasks in the current version of CryoWizard are conducted in CryoSPARC by interfacing with cryosparc-tools^14^.

The performance of CryoWizard were evaluated with six datasets downloaded from the Electron Microscopy Public Image Archive (EMPIAR): EMPIAR-10556, EMPIAR-11292, EMPIAR-10405, EMPIAR-10217, EMPIAR-10250, and EMPIAR-10876 (Fig. 6b-g). These datasets were not used in model training. The final refined maps, automatically resolved by the pipeline without performing any postprocessing, local or CTF refinement, or Bayesian polishing, achieved resolutions of 2.88 Å, 2.62 Å, 3.24 Å, 2.39 Å, 3.58 Å, and 3.01 Å, respectively. These resolutions are sufficient for accurate model building. The corresponding models of these datasets were downloaded, which align well with the resolved density maps, confirming their correctness.

### CryoWizard addresses the preferred orientation problem

The preferred orientation problem in many cryo-EM samples arises when most particles adopt a limited number of dominant poses, often due to surface charges or other unknown factors^25–27,45^. We have previously shown that Cryo-IEF can effectively cluster particles by their poses. This led us to hypothesize that certain particle classes identified by Cryo-IEF may contain particles less affected by preferred orientation, which could be used to generate isotropic maps with uniform resolutions in all directions. Therefore, we implemented an additional clustering module within CryoWizard which is triggered by the presence of preferred orientation problem (Fig. 6a and Extended Data Fig. 3).

We tested CryoWizard’s capability to address the preferred orientation problem on two datasets: EMPIAR-10217 and EMPIAR-10096, which exhibited highly imbalanced pose distributions with only a few dominant poses, as shown in the 2D classification results (Extended Data Figs. 4 and 5). Conical FSC Area Ratio (cFAR) score, which ranges from 0 to 1, evaluates the severity of the preferred orientation problem in cryo-EM maps—a lower score indicates a more severe issue^45^. Routine processing of EMPIAR-10217 and EMPIAR-10096 datasets with manually selected particles resulted in maps severely impacted by preferred orientation, with cFAR scores of 0.01 and 0.03, respectively (Fig. 7b,d and Extended Data Figs. 4 and 5). However, by clustering the Cryo-IEF-extracted particle features of EMPIAR-10217 using K-Means++ algorithm, we identified a particle class that produced a map with similar resolutions in all directions, achieving a cFAR score of 0.75 (Fig. 7a). Using this map as a template, the final refined map was exempt from preferred orientation effects and achieved a resolution of 2.37 Å (Fig. 7c). Similarly, the CryoWizard pipeline effectively addressed the preferred orientation issue in EMPIAR-10096 dataset (Fig. 7e).

**Fig. 7.**
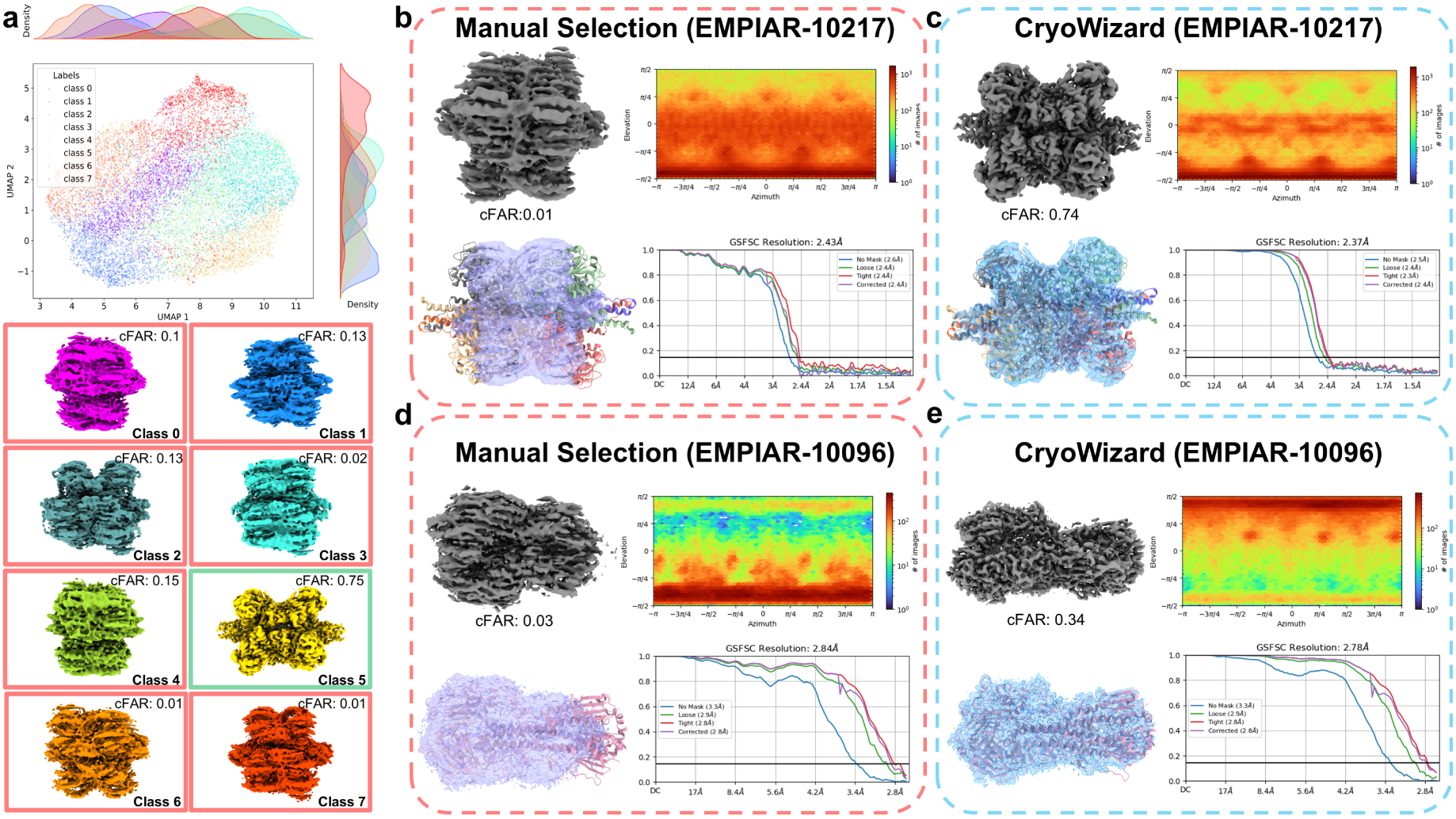
CryoWizard effectively addresses the preferred orientation problem. (**a**) For particle datasets severely afflicted by the preferred orientation problem, CryoWizard incorporates an additional module which clusters the Cryo-IEF-extracted particle features using the K-Means++ algorithm. The clustered particles from the genuine dataset EMPIAR-10217 are color-coded and visualized with UMAP. Examination of the reconstructed maps from the eight classes of particles reveals that one class (Class 5) is not affected by preferred orientation, providing a correct template for subsequent map refinement. The conical FSC Area Ratio (cFAR) scores, which indicate the severity of the preferred orientation problem, are displayed in the top right corner of each map. (**b**)(**c**) Final refinement maps reconstructed using manually selected particles (**b**) and the CryoWizard pipeline (**c**) from the EMPIAR-10217 dataset. (**d**)(**e**) Final refinement maps reconstructed using manually selected particles (**d**) and the CryoWizard pipeline (**e**) from the EMPIAR-10096 dataset. The details of manual selected particles for EMPIAR-10217 and EMPIAR-10096 datasets are shown in Extended Data Figs. 4 and 5, respectively.

## Discussion

In this study, we introduced Cryo-IEF, the first pre-trained foundation model specifically designed for cryo-EM particle image processing. Utilizing a large-scale, unsupervised model trained on approximately 65 million images, Cryo-IEF demonstrates the transformative potential of deep learning in cryo-EM data processing. Despite its unsupervised pre-training, Cryo-IEF effectively distinguishes particles based on structural variations, pose angles, and particle quality, achieving high k-NN accuracies and demonstrating three-dimensional awareness.

The development of CryoRanker, a fine-tuned variant of Cryo-IEF, further enhances the model’s performance in assessing particle quality. CryoRanker ranks particles individually, offering a significant improvement over traditional methods. Building on these capabilities, we developed CryoWizard, an automated pipeline that integrates CryoRanker to eliminate the need for manual oversight in cryo-EM data processing. CryoWizard effectively automates homogeneous structural reconstruction and demonstrates broad applicability by achieving high-resolution reconstructions across multiple datasets. By reducing consumed computational resources and manual intervention, CryoWizard improves both the efficiency and robustness of cryo-EM structural determination, significantly streamlining the processing pipeline.

CryoWizard also addresses the preferred orientation problem by identifying particle classes less affected by dominant poses. The pipeline significantly improves map isotropy, as demonstrated in datasets like EMPIAR-10217 and EMPIAR-10096. By achieving isotropic resolutions, CryoWizard provides a robust solution to a longstanding challenge in cryo-EM data processing.

Despite these advancements, our study has some limitations. First, although Cryo-IEF was trained on an extensive dataset encompassing over a hundred different biological macromolecules, samples of poor quality or low molecular weights (< 100 kDa) are underrepresented. Consequently, the model’s performance deteriorates on such samples. Expanding the training dataset to include more of these underrepresented samples will help alleviate this issue. Second, the current version of CryoWizard is designed solely for homogeneous structural reconstruction. We plan to incorporate Cryo-IEF’s classification capabilities to handle multiple structures or conformations, thus broadening its applicability and increasing the pipeline’s throughput. Third, we aim to optimize CryoWizard further to enhance its efficiency and robustness, and to explore the integration of additional tools to streamline the cryo-EM workflow.

Overall, Cryo-IEF and its applications exemplify the potential of integrating advanced AI models into cryo-EM workflows. They represent a substantial advancement in rendering cryo-EM technology more accessible, efficient and robust. By reducing consumed computational resources and manual intervention, CryoWizard has the potential to accelerate discoveries in structural biology and extend the applicability of cryo-EM across diverse research domains in biology and beyond.

## Methods

### Datasets for pre-training and fine-tuning

The datasets for pre-training and fine-tuning were gathered from various sources, including the EMPIAR database^37^, cryoPPP^38^, and in-house experiments (Extended Data Fig. 7). For most datasets, we downloaded the movies or micrographs and imported them into CryoSPARC^14^. In CryoSPARC, we performed motion correction (Patch Motion Correction; this step is skipped for micrograph data), CTF estimation (CTFFIND4^52^), particle picking (Blob Picker), and particle extraction using the tools provided by CryoSPARC (version 4.6.0). The extracted particle images were subsequently added to the pre-training dataset. For the fine-tuning dataset, we conducted particle clustering using the 2D Classification function in CryoSPARC. Based on the clarity of the 2D class averages, the cryo-EM particle images were manually assigned quality scores, ranging from 0 (pure noise or ice) to 1 (clear structural features) (Extended Data Fig. 1). Particles in the same class were assigned the same quality score. For datasets containing extracted particle images, data preprocessing step was skipped. These images were directly added to the pre-training dataset and imported into CryoSPARC for 2D classification and manually assigned quality scores before being added to the fine-tuning dataset.

In total, we gathered 65,310,474 cryo-EM particle images for pre-training the cryo-IEF model and 42,388,319 quality-labeled cryo-EM particle images for fine-tuning CryoRanker. The fine-tuning dataset is a subset of the pre-training dataset due to the exclusion of datasets with poor clustering quality (Extended Data Fig. 8) and the effort to balance the number of high- and low-quality particles.

### EMPIAR datasets

The EMPIAR IDs used for pre-training and fine-tuning are summarized (Extended Data Table 1). For each EMPIAR ID marked with an asterisk, we were able to download the particle images directly and imported them into CryoSPARC for 2D classification and assignment of quality scores. The downloaded particle images were incorporated into the pre-training dataset, while the manually-labeled particle images were included in the fine-tuning dataset. For the other EMPIAR IDs, we downloaded the movies or micrographs and imported them into CryoSPARC, where we processed the data as described above. The extracted particle images were subsequently added to the pre-training dataset, while the particle images with manually labeled quality scores were included in the fine-tuning dataset.

### CryoPPP datasets

CryoPPP comprises 34 representative protein datasets selected from EMPIAR. We downloaded all 34 datasets provided by the CryoPPP database. The downloaded particle images were imported into CryoSPARC, where we performed 2D classification. Five datasets could not be imported successfully and were therefore excluded. All 29 successfully imported datasets are listed in Extended Data Table 2. The quality of the particle images was evaluated manually based on the 2D averages. All 29 successfully imported particle image datasets were added to our pre-training dataset for cryo-IEF, while 25 datasets with manually labeled quality scores were included for fine-tuning of CryoRanker. Five datasets, including EMPIAR-10061, EMPIAR-10345, EMPIAR-10387, EMPIAR-10590, and EMPIAR-10947, were excluded from fine-tuning due to poor averages quality. Based on the 2D classification averages (Extended Data Fig. 8), the good particles could not be distinguished manually, preventing them from providing reliable scores.

### In-house datasets

We gathered 50 cryo-EM image datasets from in-house experiments that are not available in public databases such as EMPIAR. Similar to the EMPIAR datasets, these datasets consist of movies or micrographs and were processed using the tools provided by CryoSPARC. The processing pipeline mirrors that of the EMPIAR datasets. The extracted particle images were added to the pre-training dataset for Cryo-IEF training, while the particle images with manually labeled quality scores were included in the fine-tuning dataset for CryoRanker training.

### Datasets for evaluation

#### Datasets for Cryo-IEF pre-training evaluation

We downloaded 12 distinct biological particle density maps reconstructed using single-particle cryo-electron microscopy (cryo-EM) from the EMDB database^53^. These density maps were classified into four categories based on their overall molecular weights (Table 1), which were used to generate simulated cryo-EM particle images for evaluation. For each density map, we generated 2,000,000 simulated cryo-EM particle images using the Simulate Data function in CryoSPARC. The signal-to-noise ratio was set to 0.005, while other settings remained at their defaults. Based on the ground-truth pose angles recorded in the metadata generated by CryoSPARC, the simulated cryo-EM particle images were classified into 12 classes. We sampled 2,000 images for each pose angle class. To avoid ambiguity in class division, only the 50% of particles nearest to the class center were sampled as members of the final dataset. The final dataset comprised 12 × 24,000 simulated cryo-EM particle images, labeled with pose angle class and structure type.

#### Datasets for CryoRanker evaluation

We downloaded four types of genuine cryo-EM particle images from the EMPIAR database for CryoRanker evaluation. For each type, movies or micrographs were downloaded from EMPIAR and imported into CryoSPARC. Subsequently, we performed motion correction, CTF estimation, particle picking, and particle clustering using CryoSPARC, same as described in previously. For each type, we randomly selected 80,000 particle images after particle picking. The quality of the particles was scored manually in the same manner as described before (Extended Data Fig. 1).

#### Datasets for CryoSolver evaluation

To evaluate the ability of Cryo-IEF in classifying heterogeneous structures, we constructed a dataset containing particles from four different structures. The PDB IDs are 8C9C (containing 9,076 particles, 12.75%), 8C99 (containing 44,366 particles, 62.32%), 8C93 (containing 11,975 particles, 16.82%), and 8C93 (containing 5,772 particles, 8.11%). This dataset is a reduced version of the Ribosembly dataset in CryoBench^43^. The original Ribosembly dataset contains 16 types of structures. We resampled the dataset to include only four types and excluded the other 12 types with high similarity to these four structures.

#### Datasets for CryoWizard evaluation

For the evaluation of the automated pipeline (CryoWizard), we downloaded six datasets from the EMPIAR database: EMPIAR-10556, EMPIAR-10292, EMPIAR-10405, EMPIAR-10250, EMPIAR-10876, and EMPIAR-10217 (Table 2). EMPIAR-10217 and EMPIAR-10096 contain particles with preferred orientation problem, which were used to evaluate the pipeline’s performance in addressing this issue.

#### Evaluation metrics k-NN score

We utilize the k-NN score to assess the performance of cryo-IEF in distinguishing particles based on different structures, pose angles, and quality. For each data point in the test dataset, the k-NN algorithm predicts the class by examining the *k* nearest neighbors and taking a majority vote (in our experiments, *k* is set to 1). The predicted class for the *i*-th test data point is ŷ_i_, and the actual class is *y*_*i*_. The score is calculated as follows:

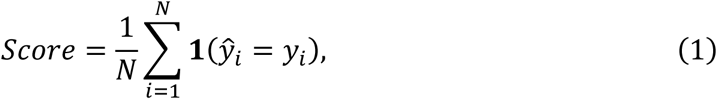

where 1(ŷ_*i*_ = *y*_*i*_) is an indicator function that equals 1 if ŷ_*i*_ = *y*_*i*_ (i.e., the prediction is correct), and 0 otherwise.

#### AUC and AP

AUC (Area Under the Curve) and AP (Average Precision) are two widely used metrics for evaluating the performance of binary classification models. In our experiments, we employ AUC and AP to evaluate the performance of the CryoRanker in distinguishing good particles from junk particles. AUC represents the area under the receiver operating characteristic (ROC) curve and can be computed as:

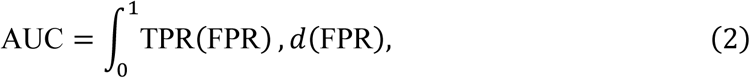

where TPR is the true positive rate and FPR is the false positive rate. AP measures the average precision at different levels of recall and is calculated as:

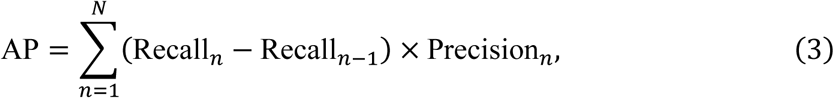

where *N* is the number of thresholds, Recall_*n*_ and Precision_*n*_ are the recall and precision at the *n*-th threshold.

#### Spearman and Pearson score

The Spearman and Pearson scores evaluate the correlation between predicted scores and labeled scores. The Pearson correlation coefficient (often denoted as *r*) is calculated as:

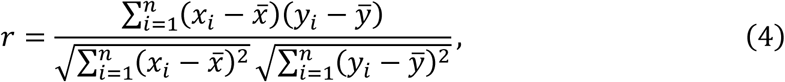

where *x^-^* is the mean of the predicted scores *x*. ŷ is the mean of the labeled scores *y*. Spearman score can be calculated as:

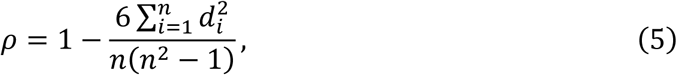

where *d*_*i*_ is the difference between the predicted score and the labeled score of the *i*-th data point.

#### Pre-training of Cryo-IEF

To enhance the training strategy for cryo-EM image data and improve stability, we implemented several modifications while generally adhering to the MOCO v3 framework.

#### Optimizer

We use the AdamW optimizer with a batch size of 2048, which empirically demonstrated good performance. The learning rate is set as lr × BatchSize/256, with the base learning rate lr set to 1 × 10^−5^. A learning rate warmup is applied during the initial 5 epochs, after which the learning rate is decayed using a cosine annealing schedule.

#### Pre-training loss

The contrastive loss function is defined as in Equation 6, where the temperature parameter τ is set to 0.5.

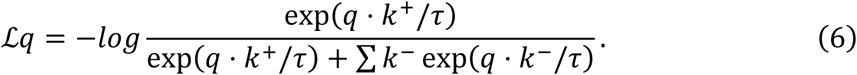

#### Data augmentation

We applied random cropping, color jittering, blurring, and solarization to the input images. Random rotation was also utilized to ensure the model’s rotation invariance. Before inputting the images into the model, they were resized to 224×224 pixels and normalized by subtracting the dataset mean and dividing by the standard deviation.

#### Model architecture

The Vision Transformer (ViT) architecture^54^ is used as the backbone of Cryo-IEF. In this paper, experiments are conducted using the ViT-B/14 backbone by default, which consists of a 12-layer transformer encoder with a hidden size of 768 and 12 attention heads.

The input patch size is 14×14, which empirically demonstrated good performance. Following MOCO v3, we replaced the Layer Normalization (LN) layer with a Batch Normalization (BN) layer in the ViT backbone’s MLP blocks (Fig. 1).

For cryo-IEF, the two encoders *f*_*q*_ and *f*_*k*_ have the same backbone and projection head architecture, with *f*_*q*_ having an additional prediction head. The projection head is a 3-layer MLP, and the prediction head is a 2-layer MLP (Extended Data Fig. 6a,b). Both have a hidden size of 4096 and an output layer size of 256. All MLPs include a Batch Normalization (BN) layer before the ReLU activation function. During training, *f*_*k*_ is updated with momentum as *f*_*k*_ = *mf*_*k*_ + (1 − *m*)*f*_*q*_, where m is set to 0.99, in alignment with MOCO v3.

#### Scale-up test

In this section, we assessed the impact of dataset size and model backbone parameters (Extended Data Fig. 6d,e). By randomly selecting different particle datasets from the original dataset, we created three additional datasets containing 25%, 50%, and 75% of the particle types, in addition to the original dataset (100% particle types). As the dataset size increased, the performance of cryo-IEF consistently improved. We also pre-trained cryo-IEF with varying backbone sizes. Notably, larger backbones did not yield improved performance, likely due to insufficient training data hindering the effective training of larger models. Thus, we posit that further increased datasets may result in additional performance gains.

#### CryoRanker: fine-tuning cryo-IEF with labeled datasets Training strategy

During the fine-tuning process, we applied a combination of cross-entropy and margin ranking loss functions to train the model. Let *D* = (*x*_*i*_, *y*_*i*_)^*N*^_i=1_ represent the dataset, where *x*_*i*_ is an input image and *y*_*i*_ is the corresponding label, onstrained such that *y*_*i*_ ∈ [0,1]. The model outputs a two-dimensional vector *p*_*i*_ = [*p*_*i*,0_, *p*_*i*,1_] for each input image *x*_*i*_, where:

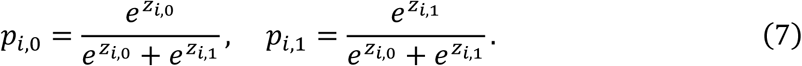

Here, *p*_*i*_ is derived from applying the softmax function to the output logits [*z*_*i*,0_, *z*_*i*,1_]. The predicted score ŷ_*i*_ of the image is defined as the probability in the first dimension: *p*_*i*,1_.

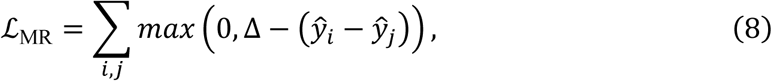

The Margin Ranking Loss is applied to the predicted scores to enforce a ranking between pairs of images. For pairs [*x*_*i*_, *x*_*j*_] where *y*_*i*_> *y*_*j*_:

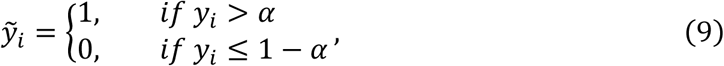

where Δ is a margin hyperparameter, empirically set to 0.2 in our experiments.

For images with labels *y*_*i*_ greater than α, we label them as 1, and for those with labels less than or equal to 1 − α, we label them as 0:

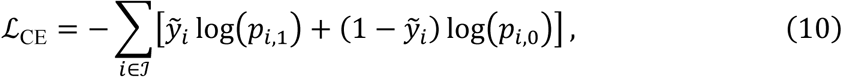

In our experiments, we empirically set α to 0.7. The Cross-Entropy Loss is then applied to these labeled elements: where ℐ is the set of indices *i* such that *y*_*i*_ > α or *y*_*i*_ ≤ 1 − α.

The final loss is the sum of the Margin Ranking Loss and the Cross-Entropy Loss:

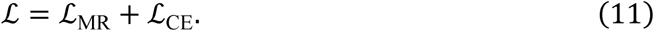

#### Ablation test

We evaluated Cryo-IEF fine-tuned with different parameters and discovered that fine-tuning the entire network yielded the best performance (Extended Data Fig. 6f), aligning with findings from previous studies in traditional computer vision^28–30,55^. We also found that the model fine-tuned with both loss functions outperformed those using either loss function alone in distinguishing particles of varying quality (Extended Data Fig. 6g). Therefore, for our experiments, we utilized CryoRanker fine-tuned with the full network and combined loss functions by default.

#### Implementation of CryoSolver

Our heterogeneous structural reconstruction experiments are conducted using the DRGN-AI framework^42^. Compared to other heterogeneous structural reconstruction methods^21,40,41^, DRGN-AI offers broader application scenarios due to its superior ab initio reconstruction performance. In our experiments, we replaced the encoder of DRGN-AI with the pre-trained Cryo-IEF model, while keeping its decoder unchanged. The new model was named CryoSolver. In CryoSolver, the decoder consists of three hidden residual layers, each with 256 units and ReLU activations. The optimization settings, including hierarchical pose search for 500,000 particles and stochastic gradient descent for 100 epochs, remain the same as in the original DRGN-AI framework. In principle, Cryo-IEF can be easily integrated into other heterogeneous structural reconstruction methods.

#### Implementation of CryoWizard

After completing the model training, quality score for input particles can be automatically assessed. However, the optimal number of particles for reconstruction is not known a priori. To address this, we have developed an automated algorithm for selecting the optimal numbers of top ranked particles based on the trained model (Extended Data Fig. 3).

#### Predicting particle scores with CryoRanker

The first stage of CryoWizard involves predicting the scores of particles using CryoRanker. The pipeline program calls CryoRanker to evaluate and record the scores of all obtained particles. Subsequently, the particles are ranked from highest to lowest scores. Assume we have a set of extracted particles *P* = *p*_1_, *p*_2_, …, *p*_*m*_, where each particle *p*_*i*_ is associated with a score *S*_*i*_. The sorted set of particles is *P*_sorted_ = *p*_1_, *p*_2_, …, *p*_*m*_, ordered such that *S*_1_≥ *S*_2_ ≥ ⋯ ≥ *S*_*m*_. During this step, extracted features from particle images are saved locally as they may be required in subsequent steps to address the preferred orientation problem.

#### Optimal subset search for initial model reconstruction

The second stage of CryoWizard involves searching for the optimal subset of particles for initial model reconstruction. Top-ranked scores are selected during this process. In the pipeline, two parallel searches are conducted with particle numbers of 50,000 and 100,000, individually. The resolution of the refined structures is evaluated, and preferred orientation analysis is performed for each structure. If both structures have a cFAR^56^ value greater than 0.15, the one with the best resolution is used as the initial structure for the following Non-uniform refinement. If neither structure meets the cFAR threshold of 0.15, an additional module for addressing the preferred orientation problem will be initiated (Fig. 6a and Extended Data Fig. 3). Further details are discussed in the section after next. In our experiments, initial model reconstruction, refinement, and preferred orientation analysis are conducted using the Ab-Initio Reconstruction, Non-uniform Refinement, and Orientation Diagnostics functions in CryoSPARC^14^, with all settings maintained at their default values.

#### Optimal subset search for resolving the final structure

The third stage of CryoWizard involves searching for the optimal number of particles for the final structural refinement. This step is analogous to the second step which is also based on the ranked scores of the particles. Let *k* represent the number of particles with scores greater than τ (where τ is empirically set to 0.4):

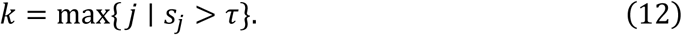

Thus, the subset of particles with scores exceeding τ is *P*_threshold_ = {*p*_1_, *p*_2_, …, *p*_*k*_}, where the particles in *P*_threshold_ is sorted by scores. *P*_threshold_ can be divided into *n* equal parts, with the *i*-th part denoted as *P*_*i*_ = {*p*_(*i*−1)⋅*k*_′_+1_, …, *p*_*i*⋅*k*_′ }, where *k*^′^. Each refinement utilizes the top *j* parts of particles, represented as ⋃^*j*^ *P*, where *j* ∈ {0,1,2, …, *n*}.

Refinement is performed based on the initial model obtained from the previous step. For all structures that meet the cFAR threshold of 0.15, the structure with the best resolution is selected as the final reconstruction. In our experiments, *n* is set to 4, as larger values are computationally expensive and yield minimal improvements. Refinement is conducted using the Non-uniform Refinement function provided by CryoSPARC^57^, with all settings maintained at their default values.

#### Implementation details of the module for addressing preferred orientation issues

The normal pipeline of CryoWizard is marked with blue arrows, while the additional pipeline for addressing the preferred orientation problem is marked with orange arrows (Fig. 6a and Extended Data Fig. 3). The preferred orientation problem is diagnosed by calculating the cFAR value of the refined structure during the initial model search step. If all searched structures have a cFAR value lower than 0.15, the preferred orientation treatment pipeline is initiated. Particles with higher scores are first selected and clustered into *n*^′^ groups (*n*^′^ is set to 8 in our experiments) using k-means++ in feature space (with the feature data saved locally in advance). The particles in each group are then selected for initial model reconstruction and refinement. For all structures with a cFAR value higher than 0.15, the one with the best resolution is used as the initial structure for the next round of refinement.

#### Integration of existing tools

Given that functions such as motion correction, CTF estimation, particle picking, and extraction can now be performed automatically, we have integrated these functions into our pipeline program. This integration enables a fully automated end-to-end process, from the movies/micrographs to the final optimized refined structure. In the diagram of the full pipeline of CryoWizard, the steps marked with dashed borders indicate these existing tools that have been incorporated into the pipeline (Extended Data Fig. 3). In our implementation, we use CryoSPARC (version 4.6.0) as the primary platform for these functions, and the pipeline program is implemented in Python by interfacing with cryosparc-tools, an open-source Python library that enables scripting access to the CryoSPARC cryo-EM software package. Theoretically, this pipeline program can be used with any cryo-EM software that provides equivalent functions.

#### Computational resources

For pre-training, we utilized four NVIDIA A100 GPUs, each with 80 GB of memory, for a total duration of 113.5 hours. For fine-tuning, we employed two NVIDIA A40 GPUs, each with 40 GB of memory, for 28.5 hours. In the heterogeneous structural reconstruction experiments, we used four NVIDIA A100 GPUs, each with 40 GB of memory, for 6.7 hours, processing a total of 71,189 particles. For the inference of CryoRanker, we utilized four NVIDIA V100 GPUs, each with 32 GB of memory, achieving a performance of 35 minutes per million particles.

## Data availability

Cryo-EM micrographs data and particle image data from EMPIAR are available at https://www.ebi.ac.uk/empiar/. Density maps used to generate simulated cryo-EM particle images are downloaded from EMDB, which is available at https://www.ebi.ac.uk/emdb/. Particles data from cryoPPP are available at https://calla.rnet.missouri.edu/cryoppp/. The raw data for 12 simulated and 4 genuine particle datasets can be found at https://drive.google.com/drive/folders/1wSdOlCdUD3LX2UJYiQ3BNJfmaNSm9zV_?usp=sharing and https://drive.google.com/drive/folders/1KxR7ZDeH_6dkyCzoHow2ANKAeGgfyTxd?usp=sharing, respectively. The resampled version of the CryoBench Ribosembly datasetis available at https://drive.google.com/drive/folders/1AXZ2hQwYowfQczRIDpSNjpmwqpKwK4PQ?usp=sharing. The reconstructed results obtained from CryoSolver and CryoWizard can be accessed at https://drive.google.com/drive/folders/1EKv5atcQA7n-VSTQ2_DW-qoVGr6g4C_X?usp=sharing and https://drive.google.com/drive/folders/1cr8nxCvkq5EAE17uZYGrJnYaxuKTwegr?usp=sharing, respectively.

## Code availability

The codes with introduction details are available at https://github.com/westlake-repl/Cryo-IEF, which is based on PyTorch.

## Acknowledgments

We thank the HPC Center of Westlake University for providing computational facility support and technical assistance. This work was supported by the Research Center for Industries of the Future (RCIF), Westlake University, the National Science Foundation of China (32122042 and 32071208 to H.S., U21A20427 to F.Y.), the Ministry of Science and Technology (MOST) of the People’s Republic of China (2022ZD0115100 to F.Y.), the Zhejiang Provincial Natural Science Foundation (DQ24C050001 to H.S.), and the Westlake Education Foundation (to H.S). We thank our colleagues Yigong Shi, Hongtao Yu, Peilong Lu, Dan Ma, Qi Hu, Qiang Zhou, Jianping Wu, Zhen Yan, Zhubing Shi, and Jijie Chai for generously sharing their in-house cryo-EM data. We would also like to acknowledge the use of data from EMDB and EMPIAR for training our models.

## Author information

These authors contributed equally: Yang Yan and Shiqi Fan.

## Contributions

The project was conceived and supervised by F.Y. and H.S. Y.Y. was primarily responsible for training the AI models, while S.F. mainly handled the preparation and processing of Cryo-EM data as well as the construction of the automated data processing pipeline. The initial draft of the manuscript was written by Y.Y. and subsequently revised and finalized by F.Y. and H.S. All authors reviewed and provided feedback on the manuscript.

## Corresponding authors

Correspondence to Fajie Yuan or Huaizong Shen.

## Extended Data Figures and Tables

**Extended Data Fig. 1.**
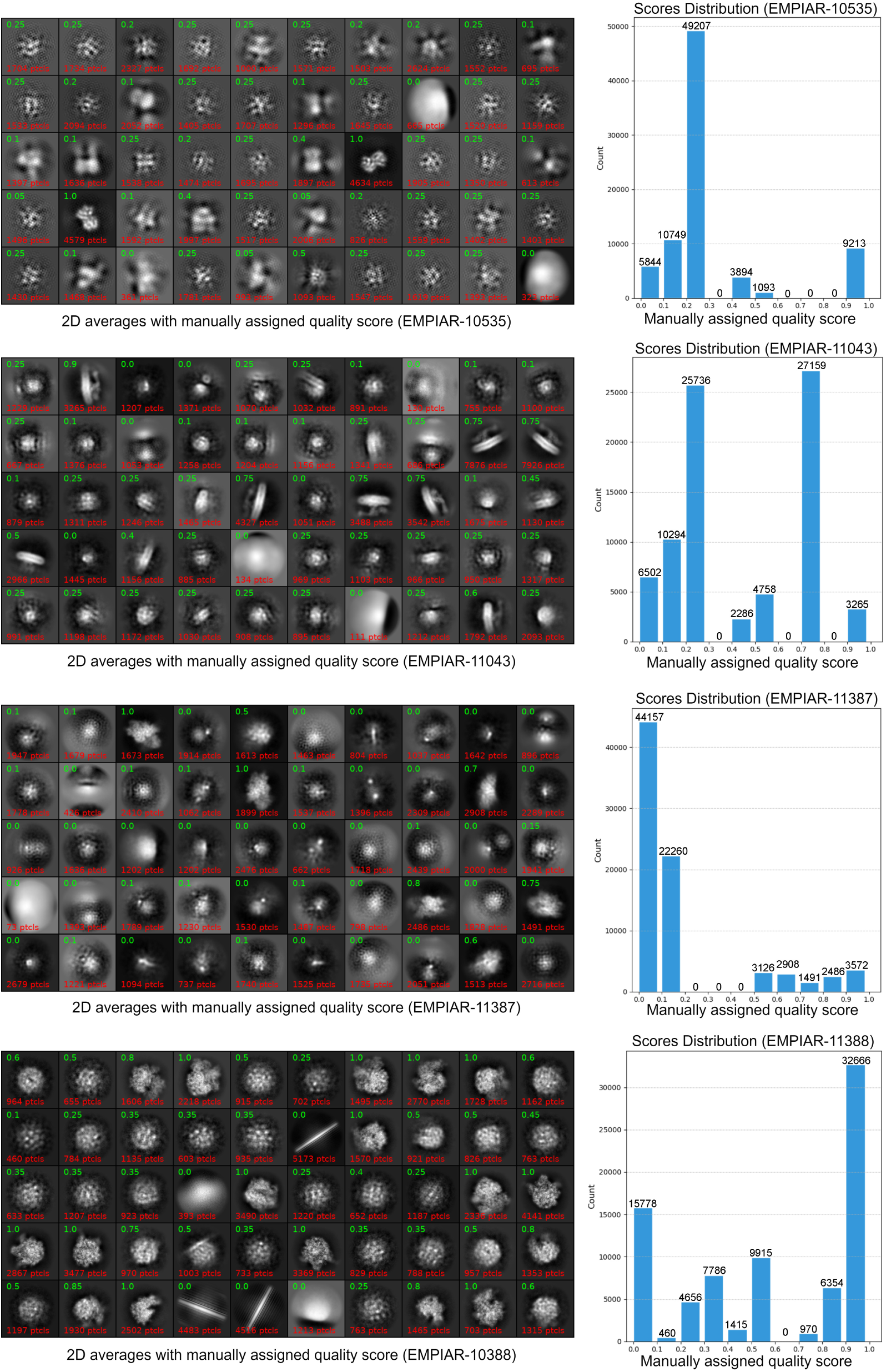
2D classification and quality score assignment of four genuine particle datasets. The cryo-EM particles of the four genuine datasets (EMPIAR-10535, 11043, 11387, and 11388) were processed using the 2D classification job in CryoSPARC and scored manually based on the quality of the 2D class averages. Distributions of the manually assigned particle scores for the four datasets are listed side by side with the 2D classification results of their respective datasets.

**Extended Data Fig. 2.**
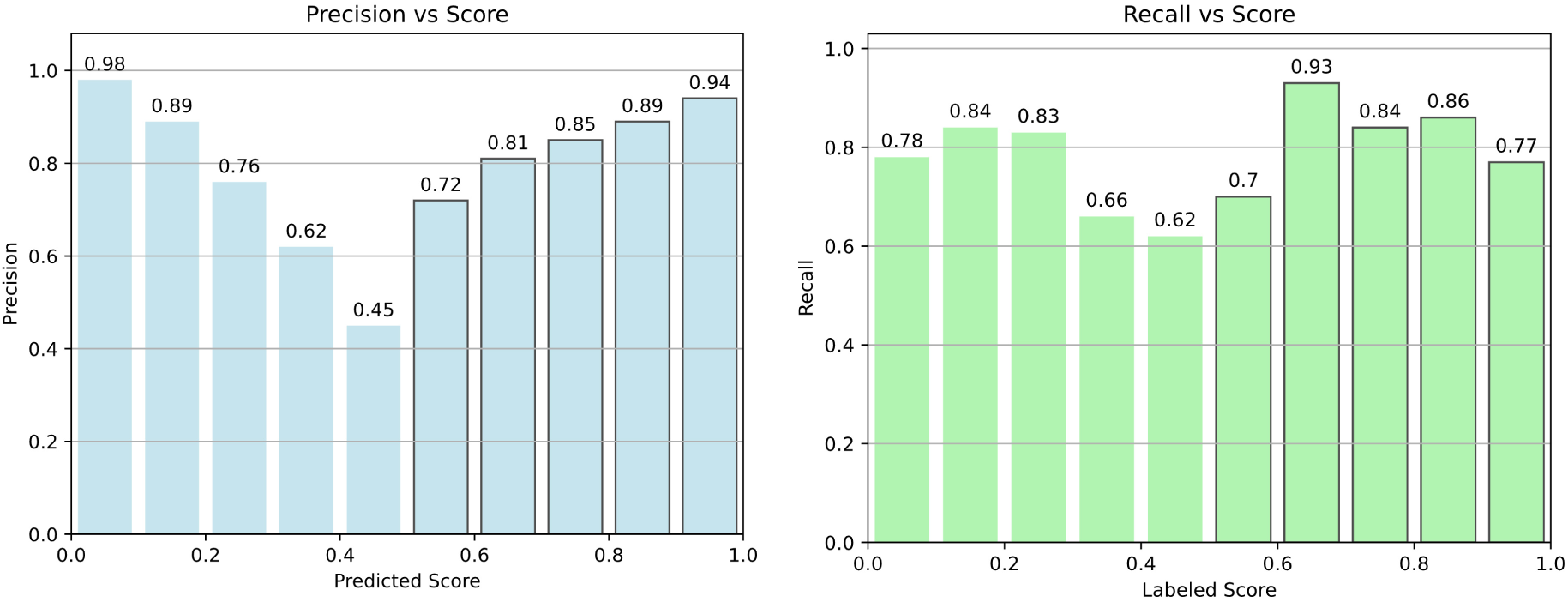
Evaluation of CryoRanker by precision and recall metrics. The precision scores in relation to the predicted particle scores (left panel) and the recall values in relation to the labeled particle scores (right panel) of the four genuine particle datasets are displayed, indicating the performance of the CryoRanker model.

**Extended Data Fig. 3.**
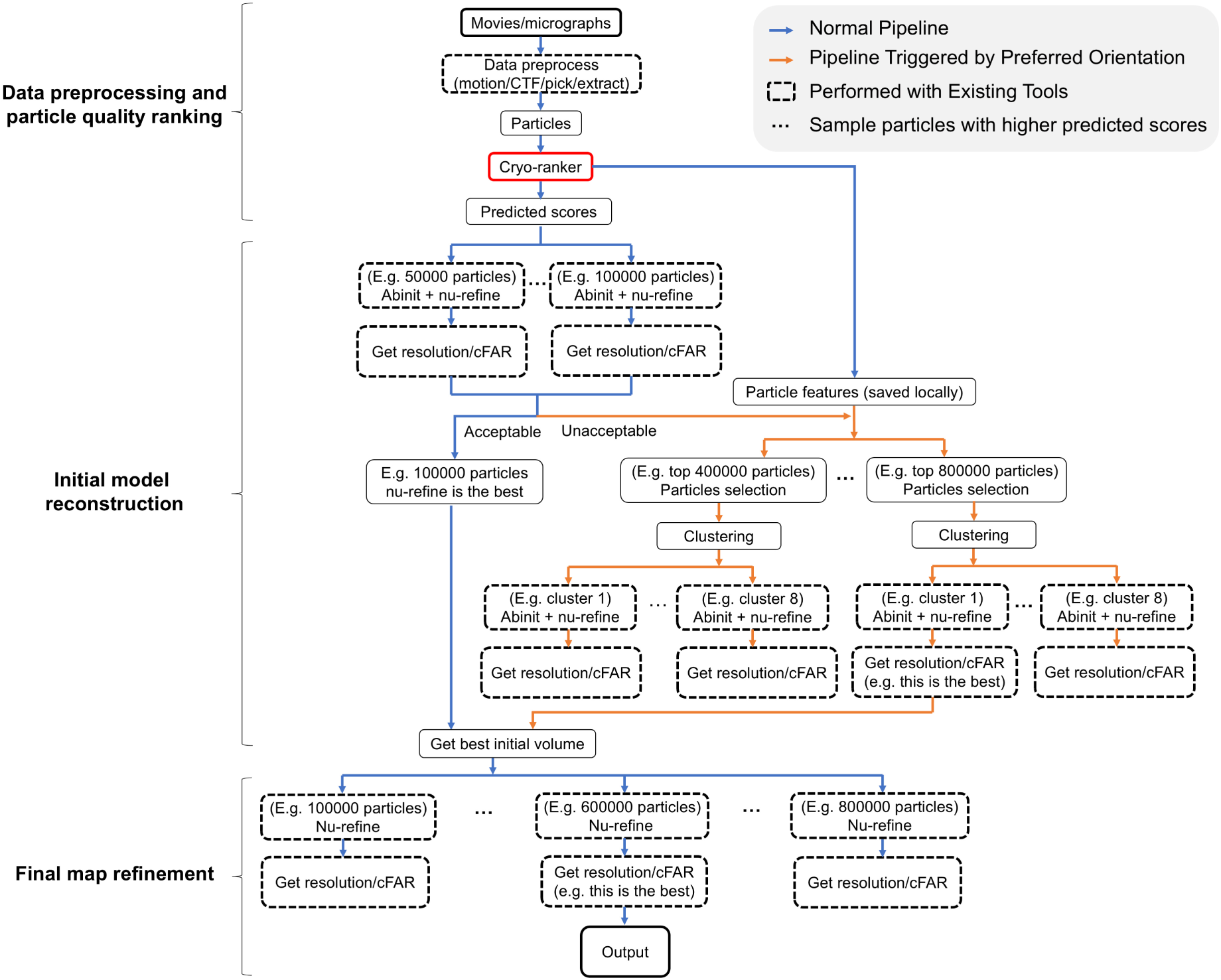
Detailed flowchart for the fully automated cryo-EM data processing pipeline, CryoWizard. The default pipeline is marked by blue arrows, while the pipeline for addressing the preferred orientation problem is marked by orange arrows. The preferred orientation problem is diagnosed by calculating the cFAR value of the refined structure during the initial model search step. Current pipeline is implemented in Python and interfaces with CryoSPARC-tools, an open-source Python library that enables scripted access to the CryoSPARC software package.

**Extended Data Fig. 4.**
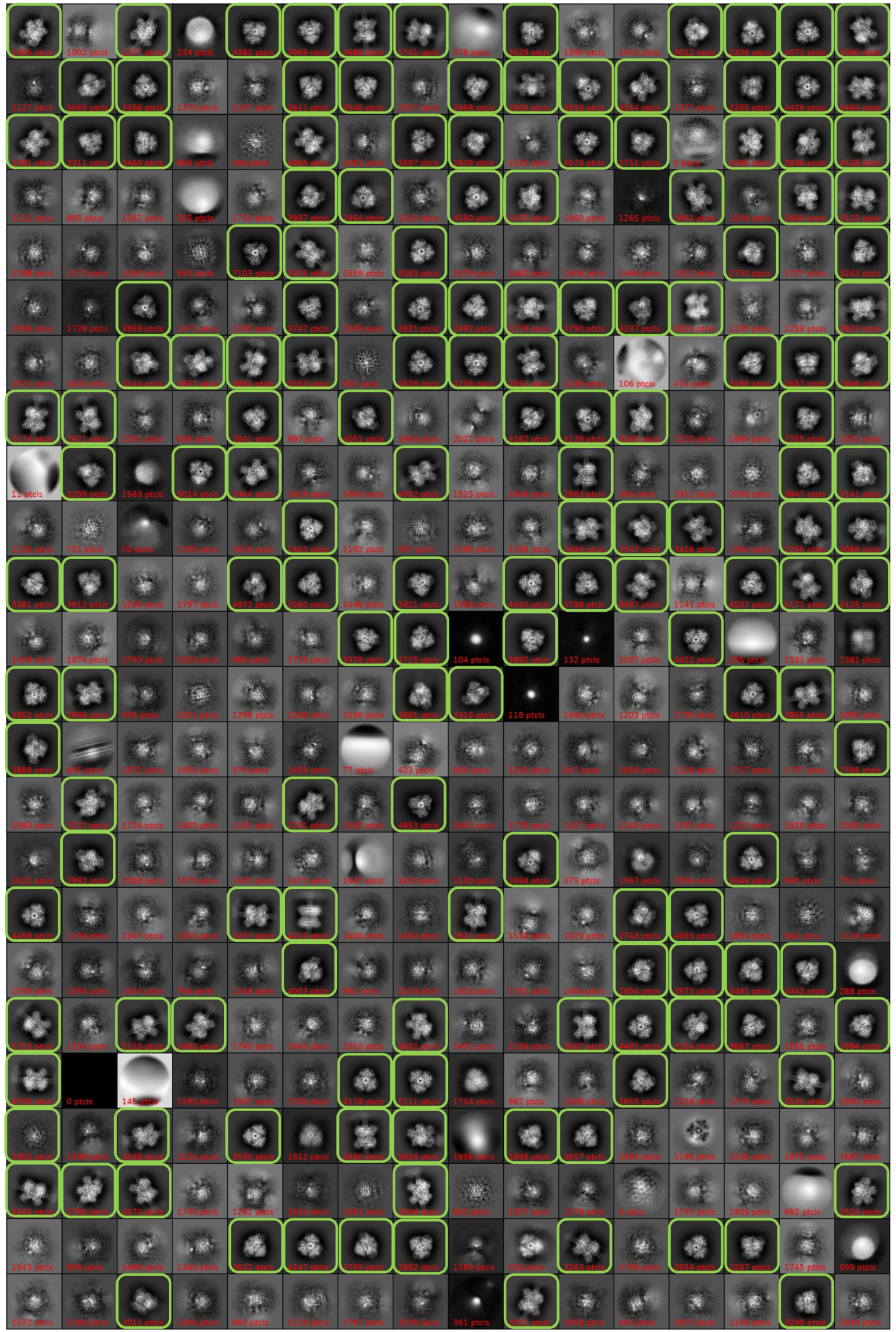
2D classification results of particles from dataset EMPIAR-10217. The cryo-EM particles of the dataset EMPIAR-10217 were processed using the 2D classification job in CryoSPARC. Manually selected classes for the following map reconstruction are indicated by the green boxes.

**Extended Data Fig. 5.**
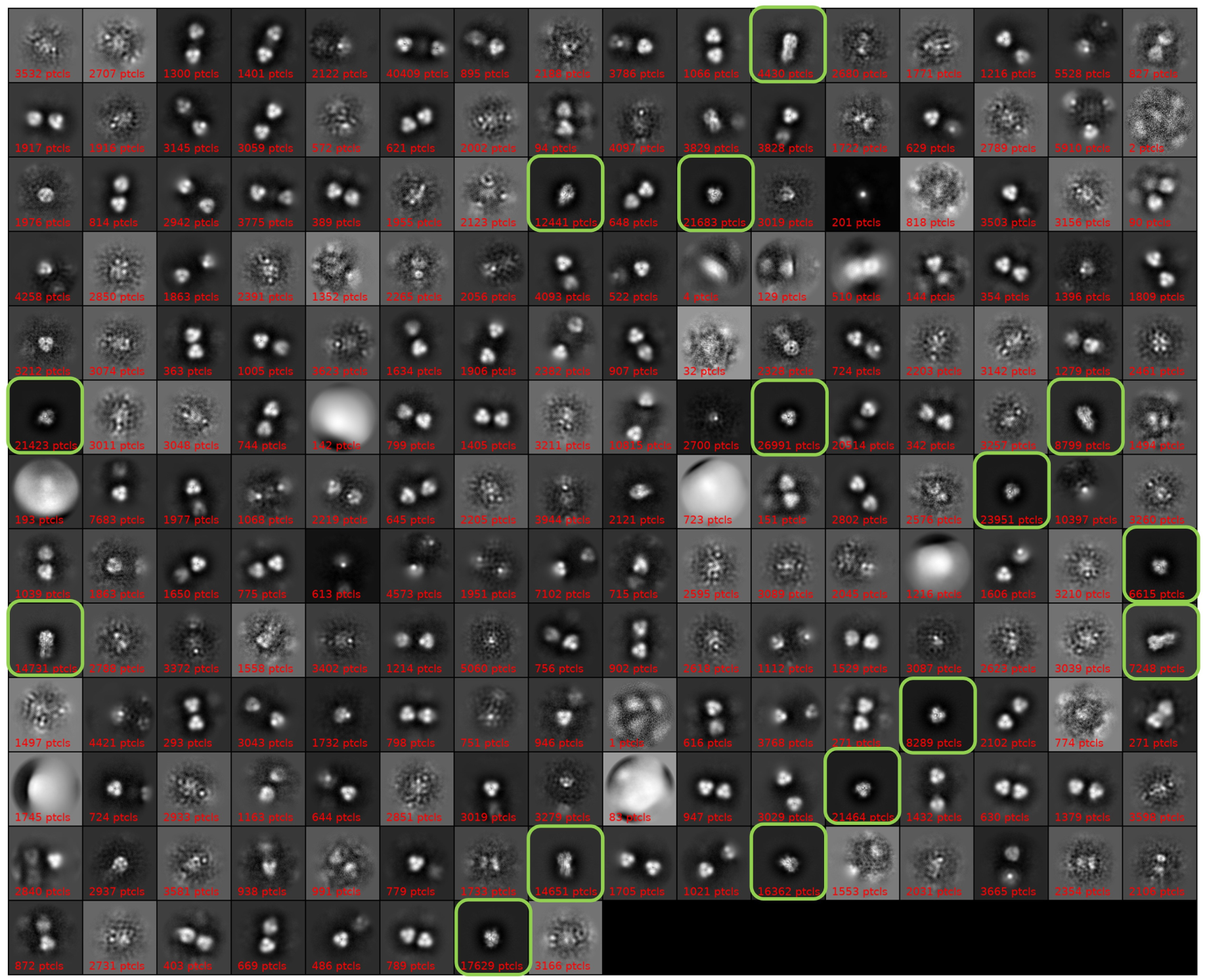
2D classification results of particles from dataset EMPIAR-10096. The cryo-EM particles of the dataset EMPIAR-10096 were processed using the 2D classification job in CryoSPARC. Manually selected classes for the following map reconstruction are indicated by the green boxes.

**Extended Data Fig. 6.**
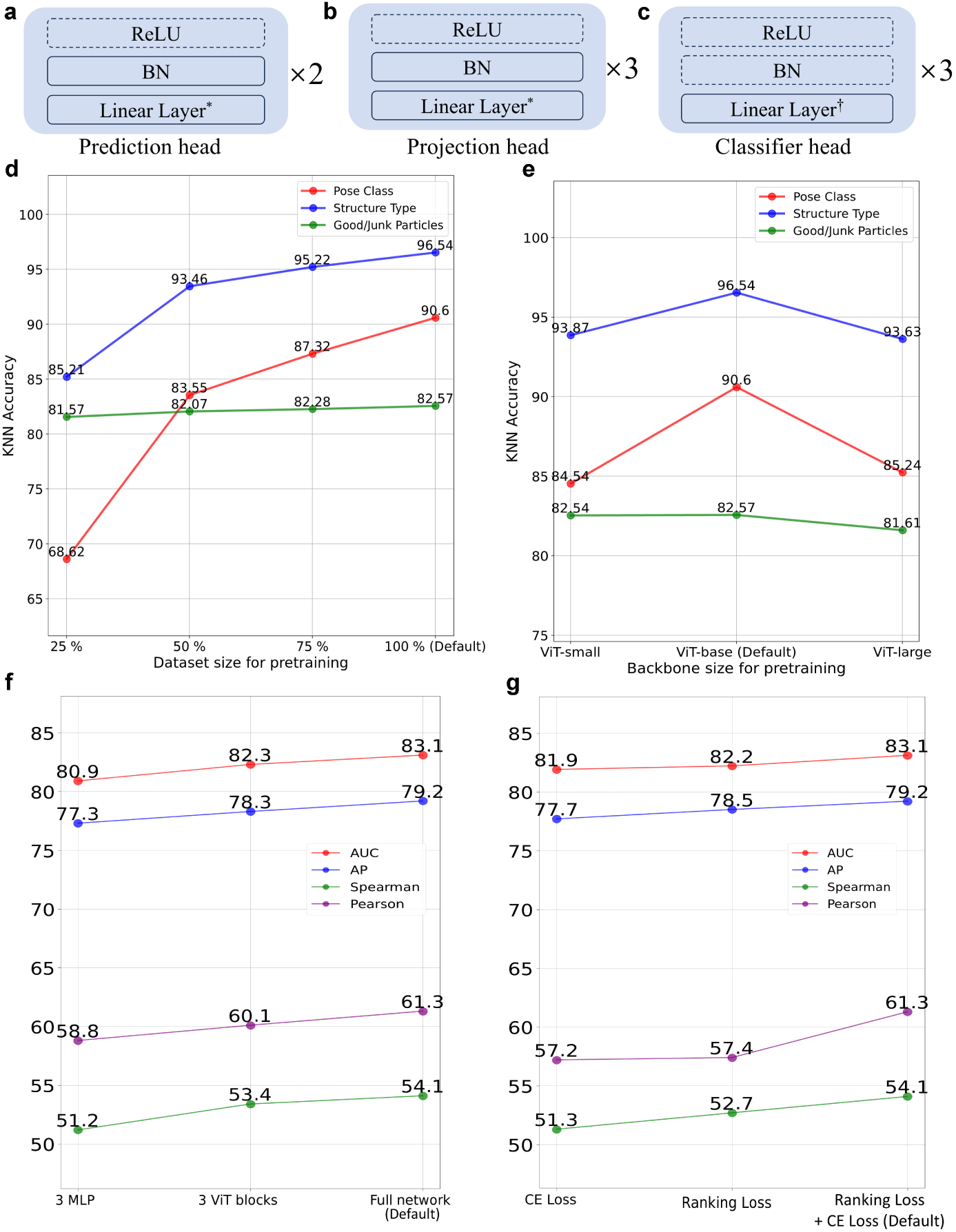
Detailed architectures and scale and ablation tests of the AI models. (**a-c**) The detailed architectures of the prediction head (**a**), the projection head (**b**), and the classifier head (**c**) in the AI models are illustrated. (**d**)(**e**) Scale test of the dataset (**d**) and backbone (**e**) sizes on the performance of the Cryo-IEF model. (**f**)(**g**) Different training parameters (**f**) and loss functions (**g**) on the fine-tuned performance of CryoRanker.

**Extended Data Fig. 7.**
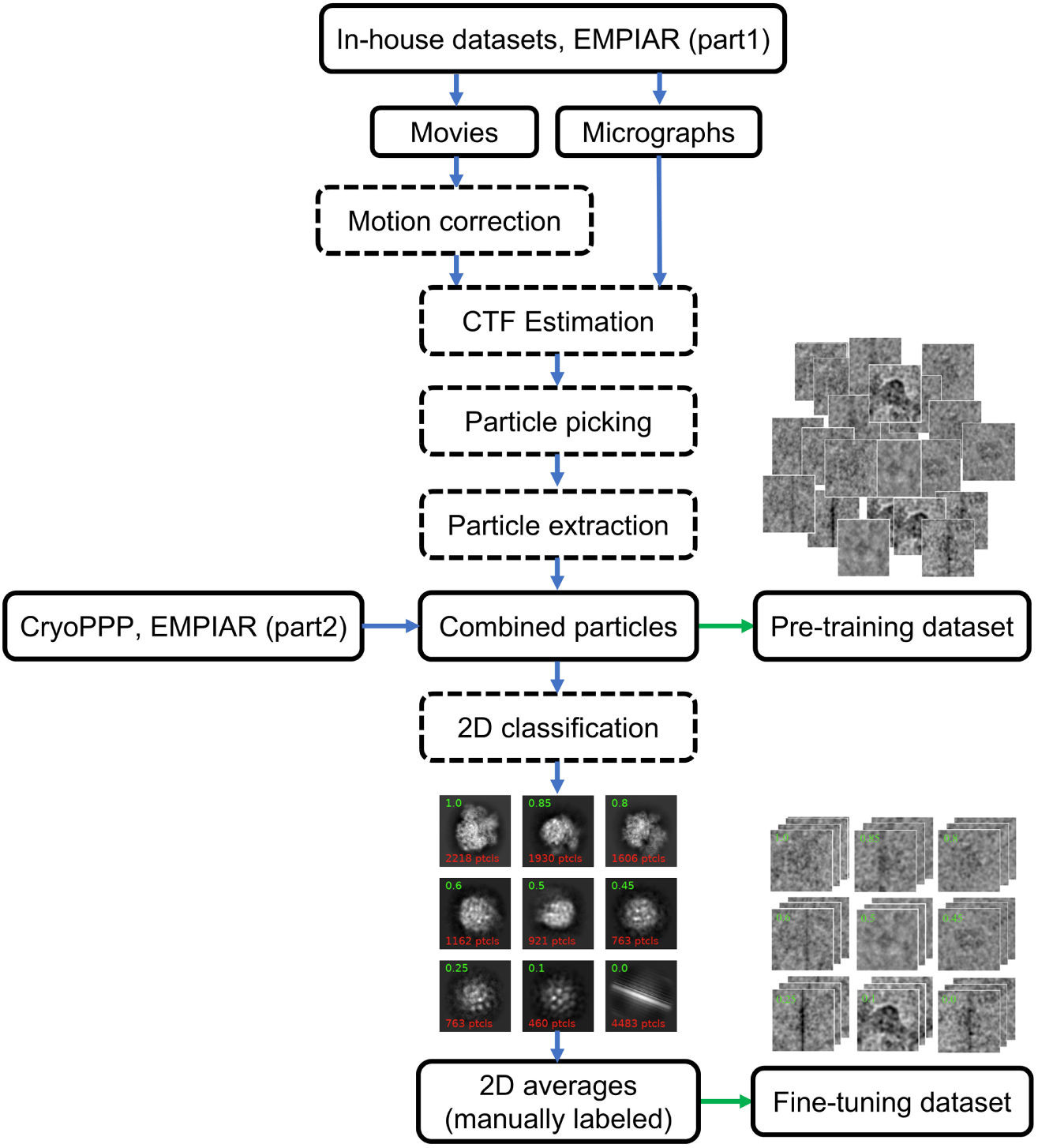
Pipeline for preparing training datasets. The pre-training dataset contains particles from various sources (EMPIAR, CryoPPP, and In-house datasets). The fine-tuning dataset is a subset of the pre-training one, where particles were processed by 2D classification in CryoSPARC and manually assigned quality scores based on the clarity of the class averages.

**Extended Data Fig. 8.**
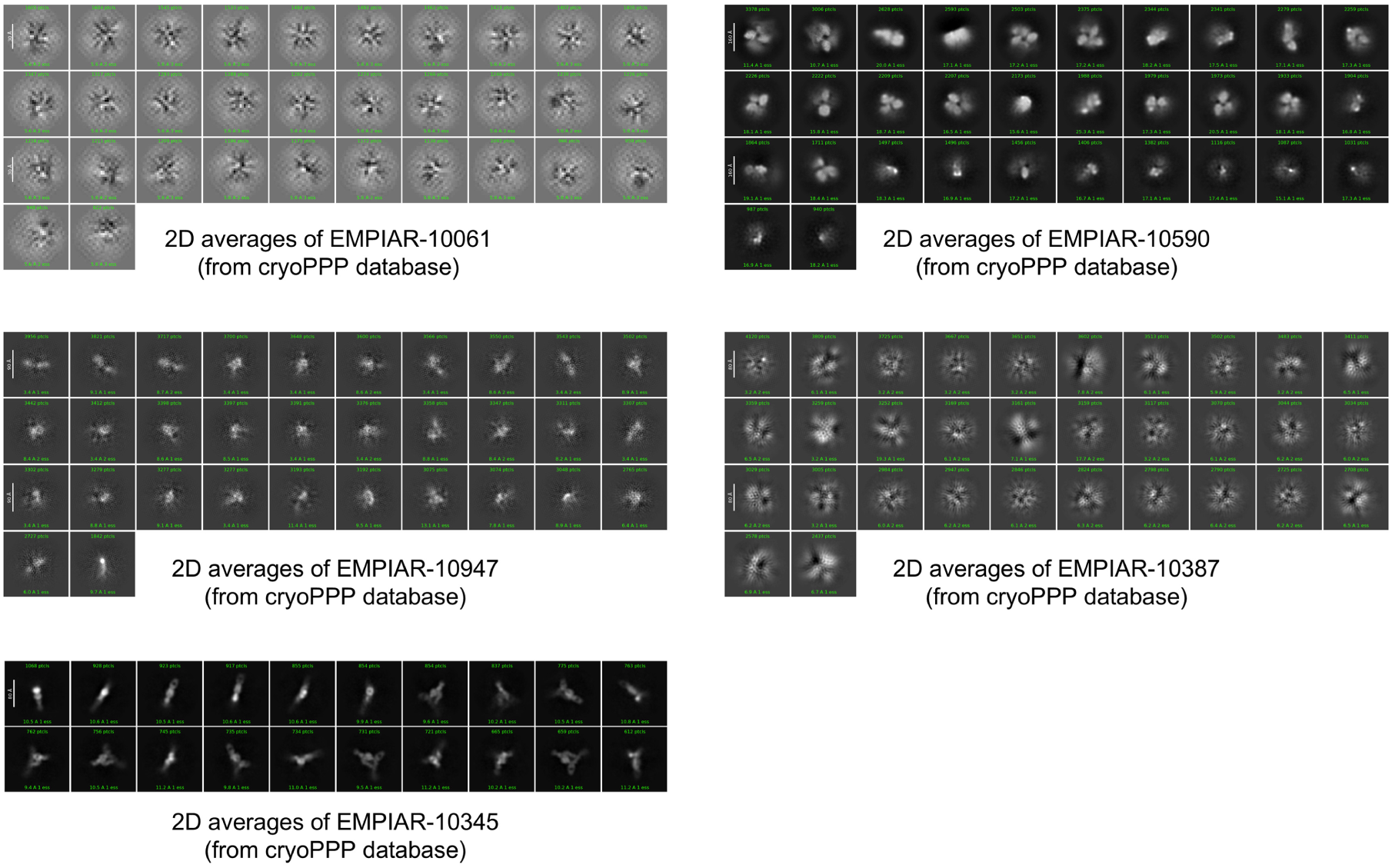
2D class averages of datasets excluded for fine-tuning. Five datasets (EMPIAR-100061, −10590, −10947, −10387, and −10345) from CryoPPP were excluded from the fine-tuning dataset due to bad qualities as visualized by 2D classification results.

---

**Extended Data Table 1.** Summary of cryo-EM data from EMPIAR.

| **EMPIAR ID** |  |  |  |  |  |  |  |
| --- | --- | --- | --- | --- | --- | --- | --- |
| 10650 | 11188 | 11265 | 11549 | 11631 | 11149 | 10381 | 10357 |
| 11738 | 11732 | 11535 | 11530 | 11202 | 10881 | 10848 | 10683 |
| 10469 | 10192 | 10969* | 10256* | 10482* | 10126* | 10127* | 11233* |
| 10255* | 10278* | 10786* | 10395* | 10970* | 10083* | 10703* | 10483* |
| 10396* | 11120* | 10254* | 10280* | 10722* |  |  |  |

\* Provided with particle images in EMPIAR.

**Extended Data Table 2.**
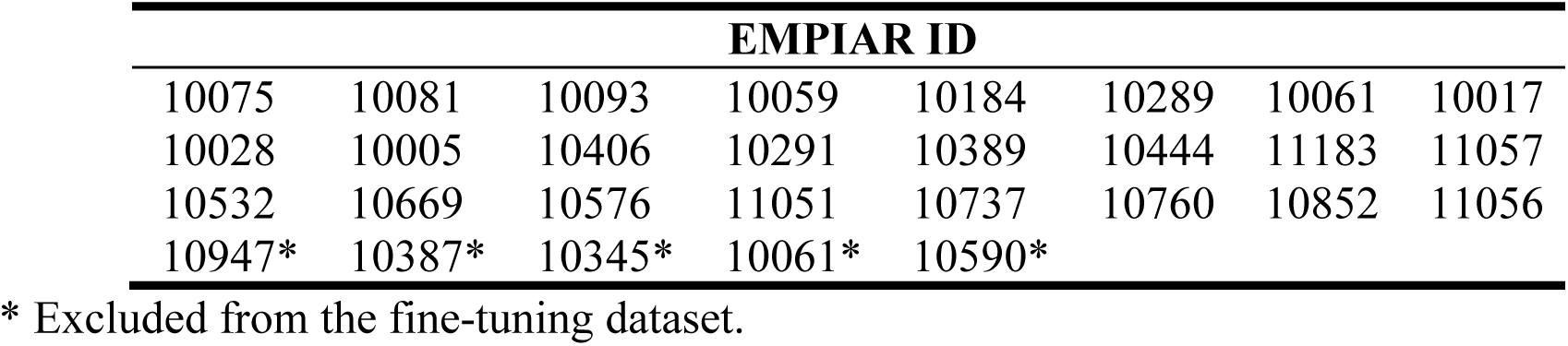
Summary of cryo-EM data from CryoPPP database.

| **EMPIAR ID** |  |  |  |  |  |  |  |
| --- | --- | --- | --- | --- | --- | --- | --- |
| 10075 | 10081 | 10093 | 10059 | 10184 | 10289 | 10061 | 10017 |
| 10028 | 10005 | 10406 | 10291 | 10389 | 10444 | 11183 | 11057 |
| 10532 | 10669 | 10576 | 11051 | 10737 | 10760 | 10852 | 11056 |
| 10947* | 10387* | 10345* | 10061* | 10590* |  |  |  |

\* Excluded from the fine-tuning dataset.

## References

1 Nogales, E. The development of cryo-EM into a mainstream structural biology technique. Nature methods 13, 24–27 (2016).

2 Cheng, Y. Single-particle cryo-EM—How did it get here and where will it go. Science 361, 876–880 (2018).

3 Bai, X.-C., McMullan, G. & Scheres, S. H. How cryo-EM is revolutionizing structural biology. Trends in biochemical sciences 40, 49–57 (2015).

4 Frank, J. Advances in the field of single-particle cryo-electron microscopy over the last decade. Nature protocols 12, 209–212 (2017).

5 Holcomb, J. et al. Protein crystallization: Eluding the bottleneck of X-ray crystallography. AIMS biophysics 4, 557 (2017).

6 Scheiner, G. The Resolution Revolution. Diabetes Self Manag 32, 28–29, 33 (2015).

7 Amunts, A. et al. Structure of the yeast mitochondrial large ribosomal subunit. Science 343, 1485–1489 (2014).

8 Liao, M., Cao, E., Julius, D. & Cheng, Y. Structure of the TRPV1 ion channel determined by electron cryo-microscopy. Nature 504, 107–112 (2013).

9 McMullan, G., Faruqi, A. & Henderson, R. Direct electron detectors. Method Enzymol 579, 1–17 (2016).

10 Zheng, S. Q. et al. MotionCor2: anisotropic correction of beam-induced motion for improved cryo-electron microscopy. Nature methods 14, 331–332 (2017).

11 Li, X. et al. Electron counting and beam-induced motion correction enable near-atomic-resolution single-particle cryo-EM. Nature methods 10, 584–590 (2013).

12 Bai, X.-c., Fernandez, I. S., McMullan, G. & Scheres, S. H. Ribosome structures to near-atomic resolution from thirty thousand cryo-EM particles. elife 2, e00461 (2013).

13 Scheres, S. H. RELION: implementation of a Bayesian approach to cryo-EM structure determination. Journal of structural biology 180, 519–530 (2012).

14 Punjani, A., Rubinstein, J. L., Fleet, D. J. & Brubaker, M. A. cryoSPARC: algorithms for rapid unsupervised cryo-EM structure determination. Nat Methods 14, 290–296, doi:10.1038/nmeth.4169 (2017).

15 Zhou, Y., Moscovich, A., Bendory, T. & Bartesaghi, A. Unsupervised particle sorting for high-resolution single-particle cryo-EM. Inverse Problems 36, 044002 (2020).

16 Zivanov, J. et al. New tools for automated high-resolution cryo-EM structure determination in RELION-3. Elife 7, doi:10.7554/eLife.42166 (2018).

17 Bepler, T. et al. Positive-unlabeled convolutional neural networks for particle picking in cryo-electron micrographs. Nat Methods 16, 1153–1160, doi:10.1038/s41592-019-0575-8 (2019).

18 Wagner, T. et al. SPHIRE-crYOLO is a fast and accurate fully automated particle picker for cryo-EM. Commun Biol 2, 218, doi:10.1038/s42003-019-0437-z (2019).

19 Kimanius, D., Dong, L., Sharov, G., Nakane, T. & Scheres, S. H. W. New tools for automated cryo-EM single-particle analysis in RELION-4.0. Biochem J 478, 4169–4185, doi:10.1042/BCJ20210708 (2021).

20 Li, Y., Cash, J. N., Tesmer, J. J. G. & Cianfrocco, M. A. High-Throughput Cryo-EM Enabled by User-Free Preprocessing Routines. Structure 28, 858–869 e853, doi:10.1016/j.str.2020.03.008 (2020).

21 Zhong, E. D., Bepler, T., Berger, B. & Davis, J. H. CryoDRGN: reconstruction of heterogeneous cryo-EM structures using neural networks. Nat Methods 18, 176–185, doi:10.1038/s41592-020-01049-4 (2021).

22 Pfab, J., Phan, N. M. & Si, D. DeepTracer for fast de novo cryo-EM protein structure modeling and special studies on CoV-related complexes. Proc Natl Acad Sci U S A 118, doi:10.1073/pnas.2017525118 (2021).

23 Jamali, K. et al. Automated model building and protein identification in cryo-EM maps. Nature 628, 450–457, doi:10.1038/s41586-024-07215-4 (2024).

24 Scheres, S. H. Processing of structurally heterogeneous cryo-EM data in RELION. Method Enzymol 579, 125–157 (2016).

25 Zhu, D. et al. Correction of preferred orientation–induced distortion in cryo– electron microscopy maps. Science Advances 10, eadn0092 (2024).

26 Zhang, H. et al. Addressing preferred orientation in single-particle cryo-EM through AI-generated auxiliary particles. bioRxiv, 2023.2009. 2026.559492 (2023).

27 Liu, Y., Fan, H., Hu, J. & Zhou, Z. H. Overcoming the preferred orientation problem in cryoEM with self-supervised deep-learning. bioRxiv, 2024.2004. 2011.588921 (2024).

28 Chen, T., Kornblith, S., Norouzi, M. & Hinton, G. in International conference on machine learning. 1597-1607 (PMLR).

29 He, K., Fan, H., Wu, Y., Xie, S. & Girshick, R. in Proceedings of the IEEE/CVF conference on computer vision and pattern recognition. 9729-9738.

30 Chen, X., Xie, S. & He, K. in Proceedings of the IEEE/CVF international conference on computer vision. 9640-9649.

31 Oquab, M., et al. Dinov2: Learning robust visual features without supervision. *arXiv preprint arXiv:2304.07193* (2023).

32 Zhou, Y. et al. A foundation model for generalizable disease detection from retinal images. Nature 622, 156–163, doi:10.1038/s41586-023-06555-x (2023).

33 Pai, S. et al. Foundation model for cancer imaging biomarkers. Nat Mach Intell 6, 354–367, doi:10.1038/s42256-024-00807-9 (2024).

34 Wang, X. et al. A pathology foundation model for cancer diagnosis and prognosis prediction. Nature, 1–9 (2024).

35 Xu, H. et al. A whole-slide foundation model for digital pathology from real-world data. Nature, 1–8 (2024).

36 Ma, C., Tan, W., He, R. & Yan, B. Pretraining a foundation model for generalizable fluorescence microscopy-based image restoration. Nat Methods 21, 1558–1567, doi:10.1038/s41592-024-02244-3 (2024).

37 Iudin, A. et al. EMPIAR: the Electron Microscopy Public Image Archive. Nucleic Acids Res 51, D1503–D1511, doi:10.1093/nar/gkac1062 (2023).

38 Dhakal, A., Gyawali, R., Wang, L. & Cheng, J. A large expert-curated cryo-EM image dataset for machine learning protein particle picking. Sci Data 10, 392, doi:10.1038/s41597-023-02280-2 (2023).

39 El Banani, M., et al. in Proceedings of the IEEE/CVF Conference on Computer Vision and Pattern Recognition. 21795–21806.

40 Zhong, E. D., Lerer, A., Davis, J. H. & Berger, B. in Proceedings of the IEEE/CVF International Conference on Computer Vision. 4066-4075.

41 Luo, Z., Ni, F., Wang, Q. & Ma, J. OPUS-DSD: deep structural disentanglement for cryo-EM single-particle analysis. Nat Methods 20, 1729–1738, doi:10.1038/s41592-023-02031-6 (2023).

42 Levy, A. et al. Revealing biomolecular structure and motion with neural ab initio cryo-EM reconstruction. *bioRxiv*, doi:10.1101/2024.05.30.596729 (2024).

43 Jeon, M., et al. CryoBench: Diverse and challenging datasets for the heterogeneity problem in cryo-EM. *arXiv preprint arXiv:2408.05526* (2024).

44 Hu, M. et al. A particle-filter framework for robust cryo-EM 3D reconstruction. Nature methods 15, 1083–1089 (2018).

45 Tan, Y. Z. et al. Addressing preferred specimen orientation in single-particle cryo-EM through tilting. Nat Methods 14, 793–796, doi:10.1038/nmeth.4347 (2017).

46 McInnes, L., Healy, J. & Melville, J. Umap: Uniform manifold approximation and projection for dimension reduction. *arXiv preprint arXiv:1802.03426* (2018).

47 Kuhlbrandt, W. Biochemistry. The resolution revolution. Science 343, 1443–1444, doi:10.1126/science.1251652 (2014).

48 Liu, Z. et al. Determination of the ribosome structure to a resolution of 2.5 Å by single-particle cryo-EM. Protein Science 26, 82–92 (2017).

49 Fan, X. et al. Single particle cryo-EM reconstruction of 52 kDa streptavidin at 3.2 Angstrom resolution. Nat. Commun. 10, 2386 (2019).

50 Baxter, W. T., Grassucci, R. A., Gao, H. & Frank, J. Determination of signal-to-noise ratios and spectral SNRs in cryo-EM low-dose imaging of molecules. Journal of structural biology 166, 126–132 (2009).

51 Palovcak, E., Asarnow, D., Campbell, M. G., Yu, Z. & Cheng, Y. Enhancing the signal-to-noise ratio and generating contrast for cryo-EM images with convolutional neural networks. IUCrJ 7, 1142–1150 (2020).

52 Rohou, A. & Grigorieff, N. CTFFIND4: Fast and accurate defocus estimation from electron micrographs. Journal of structural biology 192, 216–221 (2015).

53 Lawson, C. L. et al. EMDataBank unified data resource for 3DEM. Nucleic acids research 44, D396–D403 (2016).

54 Dosovitskiy, A. An image is worth 16×16 words: Transformers for image recognition at scale. *arXiv preprint arXiv:2010.11929* (2020).

55 Grill, J.-B. et al. Bootstrap your own latent-a new approach to self-supervised learning. Advances in neural information processing systems 33, 21271–21284 (2020).

56 Baldwin, P. R. & Lyumkis, D. Non-uniformity of projection distributions attenuates resolution in Cryo-EM. Progress in biophysics and molecular biology 150, 160–183 (2020).

57 Punjani, A., Zhang, H. & Fleet, D. J. Non-uniform refinement: adaptive regularization improves single-particle cryo-EM reconstruction. Nat Methods 17, 1214–1221, doi:10.1038/s41592-020-00990-8 (2020).

58 Pettersen, E. F. et al. UCSF ChimeraX: Structure visualization for researchers, educators, and developers. Protein science 30, 70–82 (2021).

